# A large and diverse autosomal haplotype is associated with sex-linked colour polymorphism in the guppy

**DOI:** 10.1101/2021.04.08.437888

**Authors:** Josephine R Paris, James R Whiting, Mitchel J Daniel, Joan Ferrer Obiol, Paul J Parsons, Mijke J van der Zee, Christopher W Wheat, Kimberly A Hughes, Bonnie A Fraser

## Abstract

Colour polymorphism provides a tractable trait that can be harnessed to explore the evolution of sexual selection and sexual conflict. Male colour patterns of the Trinidadian guppy (*Poecilia reticulata*) are governed by both natural and sexual selection, and are typified by extreme pattern colour variation as a result of negative frequency dependent selection. Since guppy colour patterns are often inherited faithfully from fathers to sons, it has been historically presumed that colour genes are physically linked to sex determining loci as a ‘supergene’ on the sex chromosome. Yet the actual identity and genomic location of the colour pattern genes has remained elusive. We phenotyped and genotyped four guppy ‘Iso-Y lines’, where colour was inherited along the patriline, but backcrossed into the stock population every 2 to 3 generations for 40 generations, thereby homogenising the genome at regions unrelated to colour. Using an unbiased phenotyping method to proportion colour pattern differences between and among the Iso-Y lines, we confirmed that the breeding design was successful in producing four distinct colour patterns. Our analysis of genome resequencing data of the four Iso-Y lines uncovered a surprising genetic architecture for colour pattern polymorphism. Genetic differentiation among Iso-Y lines was repeatedly associated with a large and diverse haplotype (∼5Mb) on an autosome (LG1), not the sex chromosome (LG12). Moreover, the LG1 haplotype showed elevated linkage disequilibrium and exhibited evidence of sex-specific diversity when we examined whole-genome sequencing data of the natural source population. We hypothesise that colour pattern polymorphism is driven by Y-autosome epistasis, and conclude that predictions of sexual conflict should focus on incorporating the effects of epistasis in understanding complex adaptive architectures.

## INTRODUCTION

The majority of the genome is shared by both males and females. Sexual conflict manifests when loci with opposing fitness optima arise, driving a genomic tug-of-war between the sexes. One way to resolve such conflict is to physically link sexually antagonistic (SA) loci to sex determining loci (SDL) and indeed, this framework is often used to describe the formation of sex chromosomes ^1^. However, this model neglects important factors such as pleiotropy and epistasis of sexually selected alleles, which are known to play a prominent role in the evolution of sexual dimorphism ^2,3^. Colour polymorphism provides a tractable trait for exploring the evolutionary and ecological drivers of sexual selection and conflict ^4,5^, and genome sequencing methods have hugely enhanced our ability to detect the genetic basis of colour traits ^6^. Using a unique breeding strategy designed to delineate regions of the genome related to colour, we analyse whole-genome sequencing data to uncover the genetic basis of sex-linked, sexually dimorphic colour polymorphism, lending valuable insight into the dynamics of sexual selection and sexual conflict in the Trinidadian guppy (*Poecilia reiculata*).

Guppy colour traits have fascinated biologists for a hundred years, and present an exciting system for testing predictions of sexual selection and sexual conflict. Males display a mosaic of complex and diverse colouration patterns, varying in colour, number, shape, size, and position of spots, while females are a drab and uniform tan colour ^7,8^. Guppy colour patterns exhibit high levels of standing genetic variation ^9–11^, despite evidence that mate choice and predation impose directional selection ^12–15^. Considerable evidence suggests that genetic diversity is maintained by Negative Frequency Dependent Selection (NFDS), driven by female mate preference for rare or novel morphs ^16–20^ and also frequency-dependent survival ^21,22^. Despite this great diversity in colour patterns, and our understanding of the evolutionary processes maintaining it, the underlying genetic architecture remains largely unknown.

It has long-been hypothesised that colour patterning genes and the sex determining locus (SDL) form a ‘supergene’ in the guppy ^23,24^. Early genetic studies found that at least half of guppy colour traits were Y-linked, as they are inherited completely from fathers to sons ^25,26^. Yet more recent mapping studies have highlighted the importance of X-linked and autosomal inheritance of colour traits ^27–29^. The proposed ‘Y-linked supergene’ model has motivated much of the work on guppy sex chromosome evolution, but identifying the SDL and its linked Y-specific genes has been challenging. Through multiple independent population genomic and pedigree crossing studies, it can be concluded that the Y-specific region must be small, possibly only a single gene, occurring near the distal end of chromosome twelve (LG12) ^30–32^ (but see ^33,34^). No colour or sex candidate genes have been identified in this region, and moreover, the entire LG12 is not enriched with colour genes. For example, in a QTL analysis of male colour variation, only 13% of the loci significantly associated with colour mapped to LG12 ^27^. However, intriguingly, there is evidence that the candidate Y region is highly diverse among males with many segregating male-specific variants, indicative of multiple Y haplotypes, as would be predicted under NFDS for Y-linked colour traits ^31,35^. This scenario has made it challenging to target regions of the genome responsible for colour. To date, only a handful of colour genes have been identified in colour mutants of guppies ^36,37^ and ornamental strains ^24^, and there is no evidence that these genes are located on LG12. These data suggest that colour-pattern genes are not physically linked to the SDL, but may be regulated by sex-specific loci.

We used an innovative approach to identify genomic regions associated with the highly variable, sex-linked, colour polymorphism in guppies. We phenotyped and genotyped four ‘Iso-Y’ lines, which originated from a natural population, and show strong Y-linked parental heritability in colour pattern ^19,38^. Each Iso-Y line was founded by one male showing distinct colour patterns, and was backcrossed with unrelated females in the stock population every 2 to 3 generations for 40 generations. Every generation, each line was propagated using males that closely resembled the founder of that line; thus, within each line, colour pattern was under stabilising selection. This experimental design allowed us to delineate regions of the genome related to colour pattern, as it should homogenise regions unrelated to the colour differences among lines. We then conducted an analysis of whole-genome sequencing (WGS) data from the source population, in order to examine colour-linked candidates in a natural population. Our results provide insight into the complexity of the high levels of segregating variation underpinning individual components of colour polymorphism in the guppy.

## RESULTS

### Colour patterns of Iso-Y lines are distinct across multiple dimensions of colour

Male guppies have an ultraviolet component to their colour patterns, and guppies can detect and adjust their social behavior based on ultraviolet colouration ^39^. Consequently, we used multispectral digital photography to capture human-visible and ultraviolet images of males from each Iso-Y line. We used geometric morphometrics to correct for individual differences in body size and shape among fish so that colour patterns could be measured as though they existed on identical male bodies. We then used the Colormesh pipeline ^40^ to extract colour measurements from these images at sampling locations across the body and caudal fin.

Discriminant Analysis of Principal Components (DAPC) ^41,42^ (see Materials and Methods for details) revealed that males from the different Iso-Y lines were well-separated based on colour. Discriminant functions (DF) 1, 2 and 3 accounted for 51.9%, 36.8%, and 11.4% of the variation among the Iso-Y lines, respectively (Fig. 1a), and distinguished Iso-Y9 from the other Iso-Y lines along Axis 1. Axis 2 predominately separated Iso-Y10, and axis 3 subtly differentiated the lines (Supplementary Fig. 1). Colour variation on the caudal peduncle and in the anal region were most important for differentiating the Iso-Y lines (Fig. 1b). Variation in the red colour channel associated with DF1 captures the orange colour spot observed on the anal region in Iso-Y9 (Fig. 1c).

**Figure 1.**
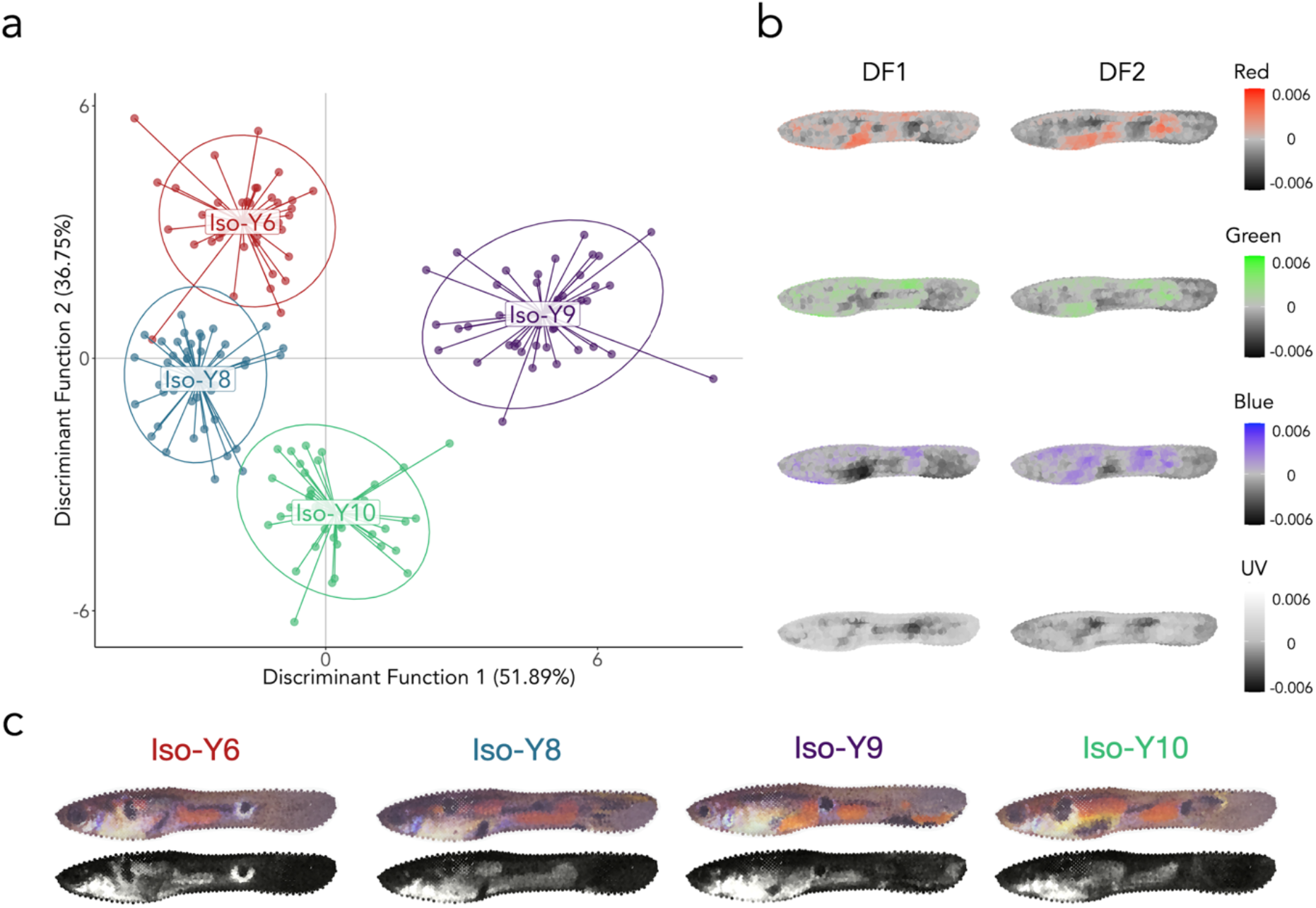
Discriminant analysis of principal components differentiating colour measurements among Iso-Y lines. (a) Scatterplot of the first 2 discriminant functions. Each point represents a male, and colour denotes the Iso-Y line. (b) Heatmaps for each colour channel depicting the correlation between colour and each sampling location and discriminant function 1 or 2. (c) Images of the male closest to each Iso-Y line’s centroid, constructed from the RGB (top) and UV (bottom) colour measures at each sampling location. UV images are false colour, with lighter grey indicating higher UV reflectance.

The Iso-Y lines also showed robust phenotypic differences when we examined mean colour measures using permutational MANOVA. Here, we reduced the dimensionality of the colour pattern data using PCA, with 17 PC’s explaining 59.3% of the total variation in colour measurements (see Supplementary Fig. 2 for PC distributions of each of the Iso-Y lines). The omnibus test indicated significant overall differences among Iso-Y lines (df = 3,169, pseudo-F = 23.66, *P* < 0.001). Post-hoc pairwise tests revealed significant differences in colour pattern among all pairs (all *P* < 0.001; see Supplementary Table 1). Based on centroid distances, the greatest phenotypic differences were between Iso-Y6 and Iso-Y10 and the smallest phenotypic differences between Iso-Y6 and Iso-Y8. Using permutational t-tests to determine whether the Iso-Y lines differ in phenotypic variance, we found that Iso-Y9 was significantly more variable (Supplementary Table 2; Supplementary Fig. 3).

### Iso-Y lines are consistently different at a large haplotype on LG1 and variable across the entire sex chromosome

Using a pooled whole genome sequencing (Pool-seq) approach and summarising all pairwise comparisons with a PCA, we were able to identify where along the genome the Iso-Y lines were consistently different (Figure 2). Pool-seq of each of the four Iso-Y lines (*n*_per line_ = 48) resulted in a final dataset of 3,995,905 SNPs. Mean pairwise *F*_ST_ between the Iso-Y lines was high overall (*F*_ST_ = 0.091). We first used pairwise *F*_ST_ values among the four Iso-Y lines, then used a multivariate approach by normalising *F*_ST_ to Z-*F*_ST_ and summarising these with PCA *F*_ST_ ^43^. The aim here was to summarise covariance of *F*_ST_ among comparisons and assess whether certain regions were consistently differentiated among the Iso-Y lines. Z-*F*_ST_ PC1 accounted for 37% of the total variance and reflected positive covariance in all pairwise *F*_ST_ comparisons (Supplementary Table 3).

**Figure 2.**
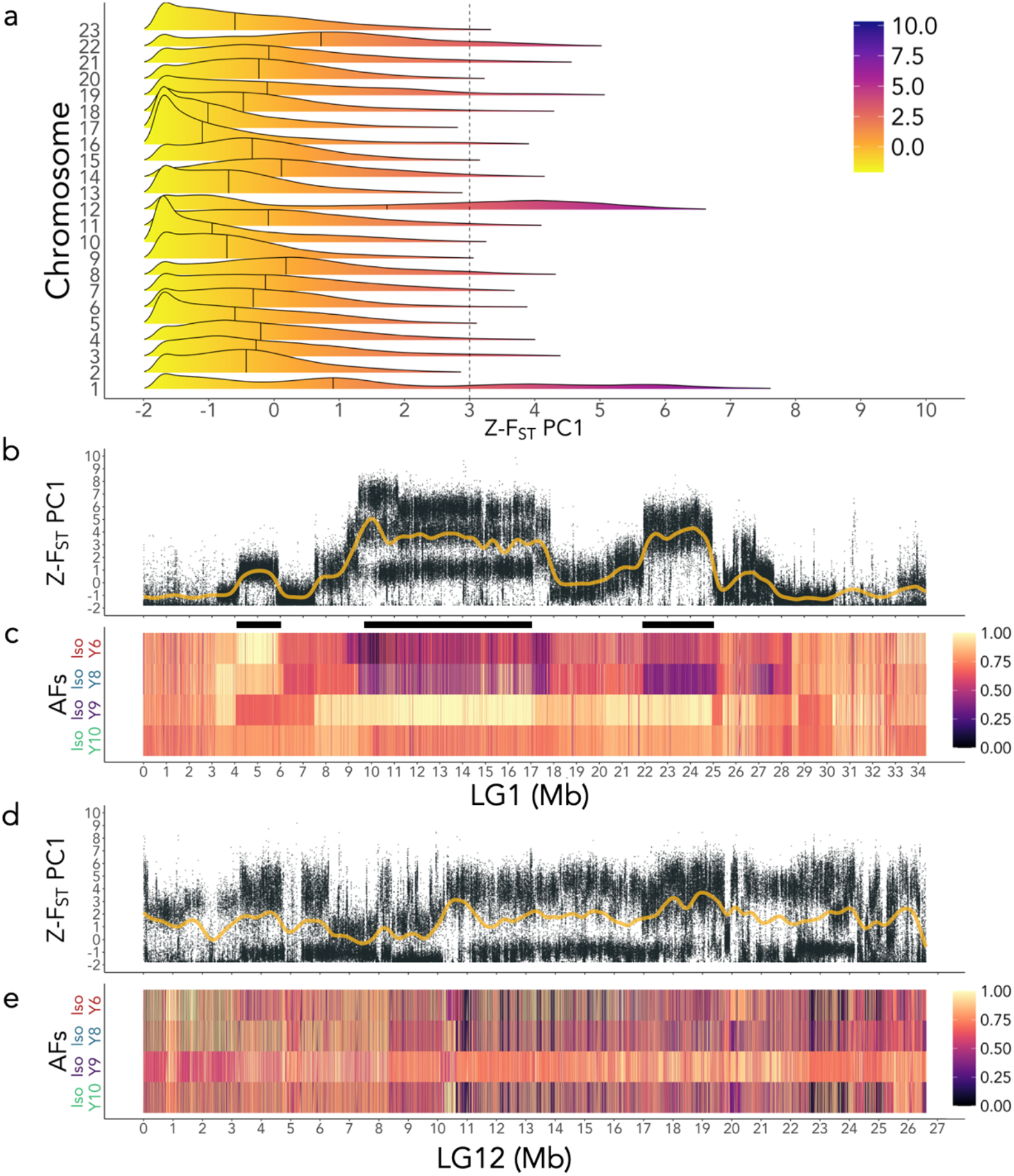
Genetic differentiation between the four Iso-Y lines: Iso-Y6; Iso-Y8; Iso-Y9; Iso-Y10. (a) Z-score PC1 *F*_ST_ density plots for each chromosome of the guppy genome (23 chromosomes). Intensity of colour scale represents higher Z-*F*_ST_ scores. Black lines within each density curve mark the median value of each chromosome. Dashed x-axis intersect marks the critical Z-score 3, α=0.05. (b) Z-*F*_ST_ PC1 for LG1; yellow line represents a smoothed spline of the data. (c) Allele frequency (AF) plots of LG1 for each Iso-Y line (Iso-Y6 - red; Iso-Y8 - blue; Iso-Y9 - purple; Iso-Y10 - green). AFs were polarised to the major allele of Iso-Y9 and changes in the AFs are represented by colour changes as depicted in the legend. On LG1, change point detection (CPD) pinpointed three regions of consistent differentiation (indicated by black rectangles): Region 1 (4-5.9 Mb); Region 2 (9.6-17 Mb); Region 3 (21.9-24.9 Mb). (d) Z-*F*_ST_ PC1 for LG12; yellow line represents a smoothed spline of the data. (e) Allele frequency (AF) plots of LG12 for each Iso-Y line. AFs were polarised to the major allele of Iso-Y9 and changes in the AFs are represented by colour changes as depicted in the legend. On LG12, change point detection (CPD) did not uncover any consistent regions of differentiation between the Iso-Y lines.

LG1 and LG12 showed excess divergence among the Iso-Y lines and were clear outliers compared to the rest of the genome (Fig. 2a). LG1 and LG12 both had the highest chromosome-wide average Z-*F*_ST_ PC1 scores (LG1 = 1.48; LG12 = 1.62) and the highest percentage of SNPs with a Z-*F*_ST_ PC1 score above a critical score of 3 (LG1 = 24%; LG12 = 29%). The remaining 47% of high-scoring SNPs were distributed across the genome, with other individual chromosomes or scaffolds accounting for <7% of high-scoring SNPs (Supplementary Fig. 4). Thus, LG1 and LG12 became our focus for investigating differentiation among the Iso-Y lines. Interestingly, the patterns of differentiation were different for these two focal chromosomes. Z-*F*_ST_ PC1 indicated three regions of high differentiation on LG1 (Fig. 2b). On LG12, these scores were elevated consistently along the entire chromosome (Fig 2d).

By performing an additional PCA on LG1, we found good agreement between the areas of differentiation amongst the Iso-Y lines. Z-*F*_ST_-LG1 PC1 (52% of the total variance) showed high positive loadings among five of the six pairwise comparisons (Iso-Y6-Iso-Y8 being the exception; Supplementary Table 4), which was also reflected in the per-SNP pairwise *F*_ST_ comparisons, where Iso-Y6 and Iso-Y8 exhibited the smallest amount of differentiation (Supplementary Fig. 5; Supplementary Table 5). The Iso-Y6-Iso-Y8 comparison loaded positively onto PC2 (17% of total variance) and Z-*F*_ST_ PC2 (and also PC3) highlighted the same regions of differentiation as PC1 (Supplementary Fig. 6).

In contrast, PC axis loadings for Z-*F*_ST_ on LG12 were different among the Iso-Y lines (Supplementary Table 6). This was also apparent in per-SNP pairwise *F*_ST_ values (Supplementary Fig. 7). All comparisons with Iso-Y9 loaded strongly onto PC1 (PC1 captured 37% of the total variance), suggesting Iso-Y9 is the most differentiated line on LG12. This was consistent with pairwise *F*_ST_, where Iso-Y9 had the highest mean pairwise *F*_ST_ > 0.2. SNPs with high Z-*F*_ST_-LG12 PC1 scores reflecting the haplotype associated with the Iso-Y9 phenotype. Z-*F*_ST_-LG12 PC2 (PC2 captured 30% of total variance) loadings were high for the remaining two comparisons with Iso-Y6. This suggests SNPs with high Z-*F*_ST_-LG12 PC2 scores reflect the haplotype associated with the Iso-Y6 phenotype. Z-*F*_ST_-LG12 PC3 reflected the remaining comparison, Iso-Y8-Iso-Y10. Z-*F*_ST_-LG12 PC2 and PC3 also reflected Iso-Y line-specific loadings (Supplementary Fig. 8). Taken together, this demonstrates that Iso-Y specific haplotypes co-occur at the same regions on LG1 whereas on LG12, the Iso-Y specific haplotypes occur in different regions.

We were able to further identify areas of consistent divergence on three regions on LG1 using change point detection (CPD) on the allele frequencies (AFs - polarised to the major allele of Iso-Y9) and Z-*F*_ST_ PC1 (Fig. 2c). Region 1 encompassed ∼1.9 Mb (coordinates: 4,079,988 - 5,984,584 bp). Region 2 comprised ∼7.4 Mb (coordinates: 9,627,619 - 17,074,870 bp). Region 3 encompassed ∼3Mb (coordinates: 21,944,840 - 24,959,750 bp). Using the same method, no regions were consistently differentiated between the Iso-Y lines on LG12 (Fig. 2e).

By examining π across LG1 (see Supplementary Text 1 for details) we found that regions of differentiation could be explained by a shared chromosomal landscape of diversity among all Iso-Y lines on LG1 (Supplementary Fig. 9; Supplementary Table 7). We found no evidence of increased coverage associated with diversity of the regions on LG1 (Supplementary Fig. 10). On LG12, we identified the previously recorded high male diversity at the putative non-recombining Y ^31^ that was shared with all Iso-Y lines (24.27 Mb) and a peak in diversity unique to Iso-Y9 at 24.28 Mb (Supplementary Fig. 11; Supplementary Table 8).

### Further examination of LG1 Region 2 reveals evidence of multiple haplotypes and colour candidate genes

In Region 2 (9.6-17 Mb), two bands of *F*_ST_ were apparent in analysis of Z-*F*_ST_-LG1 PC1, and in pairwise *F*_ST_ (Fig. 2b; Supplementary Fig. 5). To explore this further, we assessed the segregation of the AFs within each of the three identified regions in more detail (Supplementary Fig. 12). Corresponding to the double-banding of *F*_ST_, Region 2 showed unusual AF patterns within the lines. Iso-Y9 showed fixation, but the other three Iso-Y lines showed multiple bands of AFs, which taken together did not sum to 1. Additionally, in Region 3, Iso-Y6 also showed two sets of distinct bands of AFs. Assessment of the AF density distributions showed clear patterns of bimodality (trimodality in Iso-Y6) in Region 2 (Supplementary Fig. 13) and bimodality in Iso-Y6 in Region 3 (Supplementary Fig. 14). The distinct patterning corresponding to the different bands of AFs could suggest strong linkage between the SNPs segregating in each band, indicative of multiple maintained haplotypes. We reasoned that a complex haplotype structure exists, in which alleles associated with more recently derived haplotypes are nested within an older haplotype. Bimodal AF distributions parsimoniously represent AFs associated with the older (larger density peak) and younger, derived (smaller density peak) haplotypes (see Supplementary Fig. 15 for a full description).

We found several strong candidates for colour, male-specific fitness and vision in the three differentiated regions of LG1 (see Supplementary Text 2 and Supplementary Table 9). Of the 291 predicted genes within LG1 Region 2 (9.6 - 17 Mb), we identified several promising colour candidates, including *xpa*, involved in pigmentation and photosensitivity to UV light ^44^, *pcdh10a* involved in melanocyte migration, *crebbpa*, which has been identified as a candidate for plumage colouration in chickens ^45^, and *shoc2*, which causes pigmentation abnormalities ^46^, as well as five keratin genes, which have a role in pigmentation ^47^. We also identified five retinal genes (*slc24a2* ^*48*^, *stra6l* ^49^, *pnpla6* ^50^, *cabp2a* ^51^ and *nxnl1* ^52^), and three genes involved in spermatogenesis or sperm motility (*tdrd7a* ^53^, *nanos3* ^54^ and *tekt4* ^55^).

### A large variable haplotype on LG1 is maintained within the natural population

We next examined the source population using whole-genome sequencing of 26 wild-caught guppies (*n* _females_ = 16, *n* _males_ = 10, average coverage ≥13x, final dataset = 1,021,495 SNPs). Using multiple lines of evidence, we found a large haplotype on LG1 (11.1 - 15.9 Mb) segregating in the natural source population, which lies within ‘Region 2’ identified in the Iso-Y analysis (9.6 - 17 Mb). We term this area ‘Region 2-NP’ (Region 2 Natural Population) (Figure 3).

**Figure 3.**
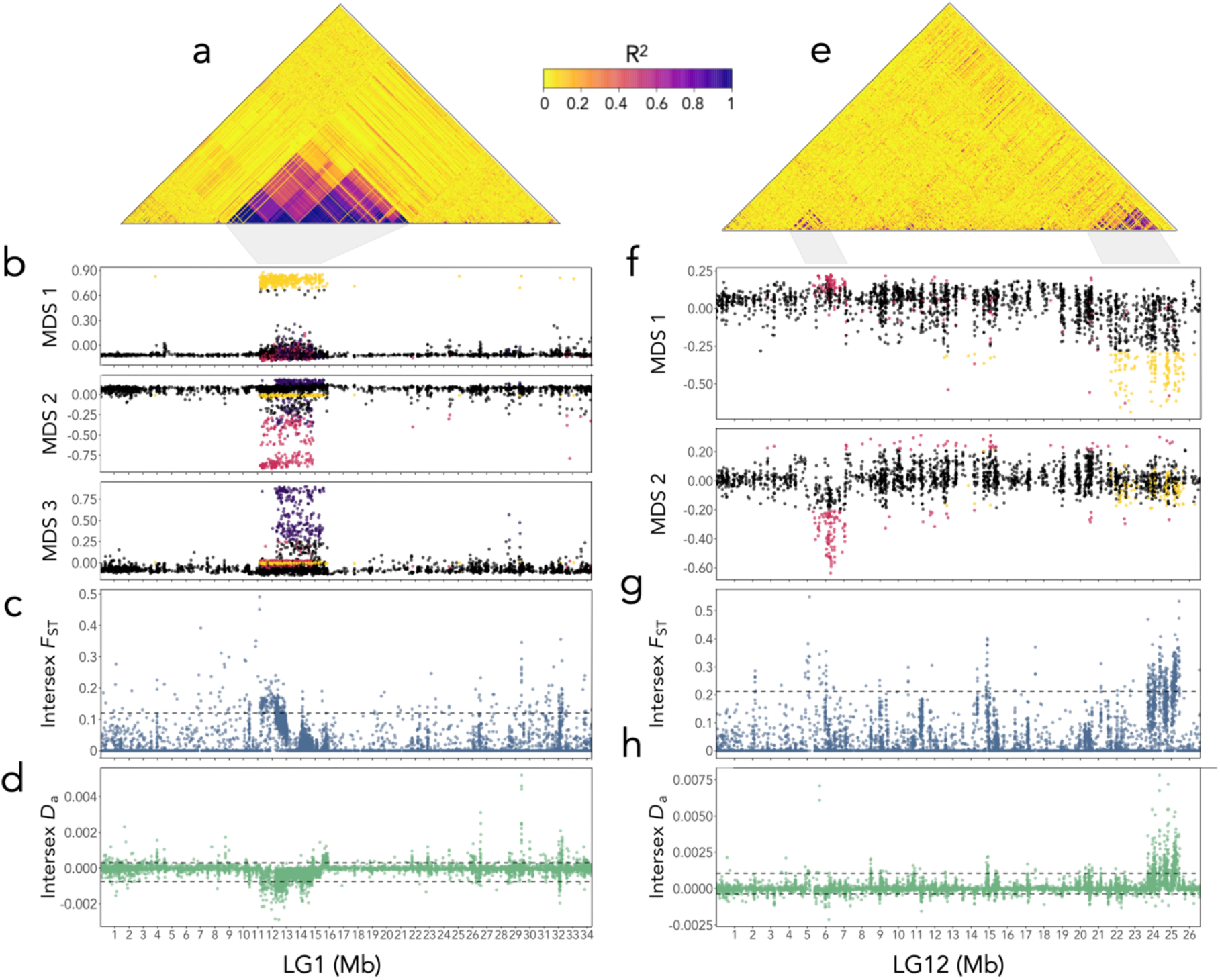
Analysis of LG1 and LG12 in the natural source population (*n* _females_ = 16, *n* _males_ = 10). (a) LG1 heatmap of patterns of linkage disequilibrium (LD) showing high LD (coordinates: 11,114,772 - 15,890,374). (b) LG1 local PCA in 10bp windows, depicting three significant MDS: MDS1 (yellow, coordinates: 11,216,906 - 15,375,083 bp); MDS2 (pink, coordinates: 11,430,012 - 14,570,259 bp); MDS3 (purple, coordinates: 12,326,328 - 15,308,055 bp). (c) Intersex *F*_ST_ across LG1. Dashed lines mark 95% quantiles. (d) Intersex *D*_a_ across LG1. Dashed lines mark 5% and 95% quantiles. (e) LG12 heatmap of patterns of linkage disequilibrium (LD) showing high LD between ∼4.6 - 6Mb and at the terminal region between ∼23.8 - 25.3 Mb. (f) LG12 local PCA in 10bp windows, depicting two significant MDS: MDS1 (yellow, coordinates: 21,913,633-25,664,533 bp); MDS2 (pink, coordinates: 5,608,703 - 7,063,435 bp). (g) Intersex *F*_ST_ across LG12. Dashed lines mark 95% quantiles. (h) Intersex *D*_a_ across LG12. Dashed lines mark 5% and 95% quantiles.

Analysis of linkage disequilibrium (LD) on LG1 revealed an area of high linkage between 11,114,772 to 15,890,374 bp, where it was apparent that several overlapping linkage blocks exist (Fig. 3a). We also analysed patterns of LD for males and females separately, and found that males exhibited at least two distinct neighbouring linkage blocks, but females showed high linkage across the entire region (Supplementary Fig. 16). We then assessed shifts in local ancestry by performing a local PCA approach ^56^, which recapitulated the identified areas of high LD (Fig. 3b), with a pattern of significant differentiation (saturated eigenvalues > 0.01) represented by three overlapping Multidimensional scales (MDS) on LG1: MDS1: 11,216,906-15,375,083 bp; MDS2: 11,430,012-14,570,259 bp; MDS3: 12,326,328-15,308,055 bp. This demonstrates that subsets of correlated SNPs within Region 2-NP exhibit ancestry relationships that deviate from those observed across the rest of the chromosome; a pattern indicative of inversions, changes in recombination, or gene density ^56^.

Intersex *F*_ST_ showed differences between the sexes within Region 2-NP (Fig. 3c); in particular, an area of high density of elevated intersex *F*_ST_ between 12.2 Mb to 13.1 Mb. We found that elevated intersex *F*_ST_ was driven by a reduction in male-specific diversity, as measured by intersex *D*_a_ (*D*_XY_ - female π; Fig. 3d). Overall, these results suggest that selection is operating differently between the sexes in our candidate region, indicating in particular, signatures of positive selection in males.

As recombination history is known to affect the maintenance of tightly linked genetic architectures ^57^, we also examined the genome features of Region 2-NP. We did not observe any differences in the proportion of GC content (Supplementary Fig. 17), nor repeat elements (Supplementary Fig. 18) in our candidate region. SNP density within Region 2-NP was considerably higher compared to the rest of the chromosome (Supplementary Fig. 19), but there was no evidence of extremes in read depth indicative of duplication or copy-number variation (Supplementary Fig. 20). Gene density showed a moderate number of genes across Region 2-NP, and a drop in gene density at ∼13.2Mb (Supplementary Fig. 21).

We performed similar analyses on LG12, confirming previously identified differences between the sexes ^30,31,58^. LD analysis showed two main regions of increased linkage; a fragmented region extending from 4.6 - 6.7 Mb, and another more clearly defined region near the terminal end of the chromosome (23.8 - 25.3 Mb) (Fig. 3e). A local PCA of LG12 identified two significant MDS axes (saturated eigenvalues > 0.01) approximately corresponding with the areas of high linkage (Fig. 3f). MDS1 outliers were observed at the terminal end of the chromosome (coordinates: 21,913,633-25,664,533 bp). MDS2 depicted an area encompassing the latter part of the high linkage block, extending past it in the latter coordinates (coordinates: 5,608,703 - 7,063,435 bp; Fig. 3e).

Overlapping with MDS2, a significant elevation in intersex *F*_ST_ was observed (Fig. 3g) and in the latter part of the region, a significant narrow peak in increased male diversity was detected at 5.67 Mb (Fig. 3h). This second region overlaps with a previously identified *D*_a_ outlier window in an analysis of six natural populations (6.94 - 6.95Mb and 7.00 - 7.01 Mb), where male-specific *D*_a_ was notably higher in high-predation (HP), compared to low-predation (LP) populations ^31^. Moreover, the LD in this region extends across 4.6 - 6 Mb which includes another previously identified male-biased candidate region (LG12: 4.8 - 5.2 Mb) ^31^. An assessment of the gene annotations within the MDS2 coordinates identified only a few candidates of interest (Supplementary Table 10): *dmgdh*, a gene which affects sperm trait variation and is part of a sex-supergene in songbirds ^59^ and *bhmt*, a folate-related gene which shows an association with skin pigmentation in humans ^60^.

It is predicted that the terminal region of LG12 contains the sex-determining locus (SDL) ^27,61,62^ and a recent multiple population genomics survey identified a sex-linked region between 24.5 and 25.4 Mb in LG12, which overlaps with the region of high LD and MDS1 region identified here ^31^. Moreover, analysis of intersex *F*_ST_ showed high differentiation between the sexes in the region (Fig. 3g), and analysis of intersex *D*_a_ revealed that this was driven by an excess of male diversity (Fig. 3h). These results are consistent with a hypothesis of multiple diverse Y-haplotypes and NFDS on the Y. Previous investigation of the gene content within the terminal region found it to be relatively gene-poor, containing multiple repeated copies of NLRP1-related genes ^31^. Our analysis of gene content for the slightly expanded region found for this single population identified a single potential colour pattern candidate (*vldlr*), which is responsible for caudal fin patterning in *Danio rerio* ^63^, and two Y candidates: *AIG1* family, and *spag16* (Supplementary Table 10).

Combining both the natural population data and our experimental Iso-Y lines, we found evidence for the maintenance of large, highly diverse haplotypes on LG1 associated with male-specific colour. In the Iso-Y lines, Region 2 encompassed multiple bands of tightly associated allele frequencies, suggesting multiple derived haplotypes. Analysis of the natural population data revealed that Region 2-NP is characterised by elevated linkage, and is defined by linkage patterns indicative of not just one, but multiple derived local ancestries. Comparing male and female diversity within the population suggests selection may operate differently between the sexes in Region 2-NP; in particular, between 12.2 Mb and 13.1 Mb, which showed high intersex *F*_ST_ and a reduction in male diversity. SV analyses using short-read and long-read data did not show support for SVs in Region 2 or Region 2-NP (Supplementary Table 11). Our prediction is that the region encompasses a large and diverse haplotype, and that within this haplotype, genetic rearrangements have resulted in several derived haplotype segments. To interrogate this further, we performed an in-depth analysis.

Individual genotypes of the Iso-Y lines and the natural population data across LG1 revealed long stretches of maintained genotype states within Region 2-NP (Fig. 4a). In the Iso-Y line data, Iso-Y9 showed distinct blocks of fixed HOM ALT genotypes, whilst the other Iso-Y line genotypes were entirely heterozygous (HET). This observation is congruous with the multiple bands of allele frequencies observed in the Iso-Y line data. Of individuals from the natural population (*n* _total_ = 26), 17 (*n* _females_ = 10, *n* _males_ = 7) shared the same overall genotype signature of HOM ALT genotype blocks as seen in Iso-Y9. Seven individuals (*n* _females_ = 4, *n* _males_ = 3) showed genotype blocks alternating between areas with the Iso-Y9 HOM ALT genotype signature, and areas of heterozygosity. Lastly, only two individuals showed a signature of extended blocks of HOM REF genotypes: NAT08 and NAT16 (*n* _females_ = 2, *n* _males_ = 0), yet HOM REF genotypes were only maintained for a proportion of the entire region.

**Figure 4.**
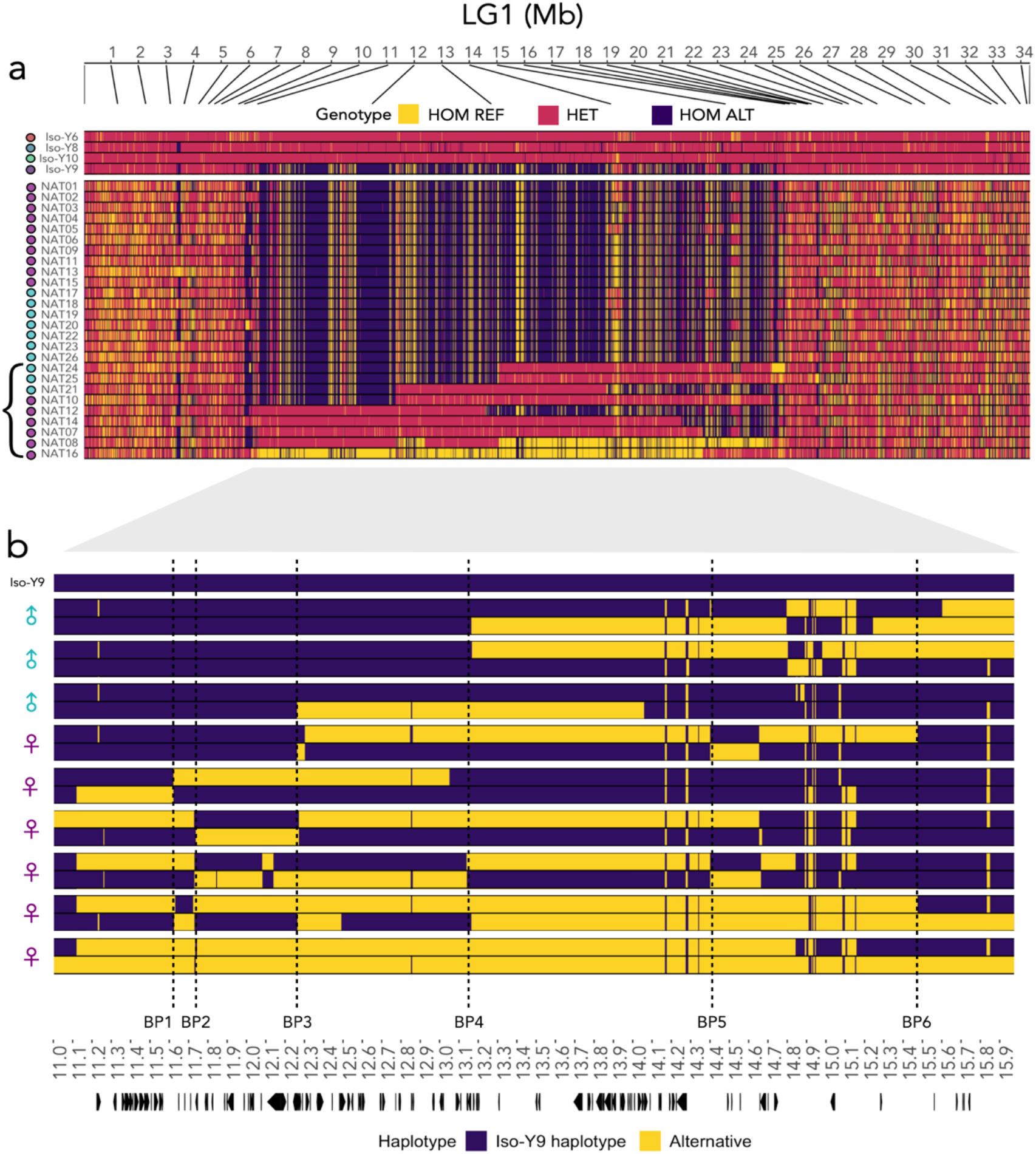
LG1 ‘Region 2’ haplotype structure. (a) Genotype plot for Iso-Y lines and natural data combined showing the genotype of each SNP depicted as homozygous reference (HOM REF: yellow), heterozygous (het: pink) or homozygous alternative (HOM ALT: purple); each individual is coloured by Iso-Y line, or by sex; females in purple, males in cyan. Bracket represents heterozygous individuals used in the haplotype analysis below. (b) Haplotype plot of phased data for natural-derived heterozygous samples (*n* _total_ = 9, *n* _males_ = 3, *n* _females_ = 6) polarised to the Iso-Y9 haplotype (purple) and alternative (yellow). Symbols next to each of the individuals represent sex: males in cyan, females in purple. Breakpoints in the haplotype are when phases switch between purple and yellow. Dashed lines mark conserved breakpoints (≥2 individuals): BP1: 11.6 Mb (*n* _females_ = 2); BP2: 11.7 Mb (*n* _females_ = 3); BP3: 12.2 Mb (*n* _females_ = 3, *n* _males_ = 1); BP4: 13.1 Mb (*n* _females_ = 2, *n* _males_ = 2); BP5: 14.4 Mb (*n* _females_ = 2); BP6: 15.4 Mb (*n* _females_ = 2). Arrows at the bottom highlight the location of gene annotations for the region.

By phasing the heterozygous individuals (*n* _total_ = 9), we found evidence of two large divergent haplotypes encompassing the entirety of Region 2-NP with subsequent recombination at conserved breakpoints (Fig. 4b). This indicates that the whole high linkage block is not one large inversion. We identified six repeated breakpoints (i.e. occurring in ; ≥ 2 individuals): BP1: 11.6 Mb (*n* _females_ = 2); BP2: 11.7 Mb (*n* _females_ = 3); BP3: 12.2 Mb (*n* _females_ = 3, *n* _males_ = 1); BP4: 13.1 Mb (*n* _females_ = 2, *n* _males_ = 2); BP5: 14.4 Mb (*n* _females_ = 2); BP6: 15.4 Mb (*n* _females_ = 2). Breakpoints were primarily located in females, except for BP3 and BP4. This difference between the sexes in the conservation of these breakpoints suggests sex-biased recombination in breaking up the overall haplotype.

These analyses allowed us to pinpoint breakpoints contributing to patterns of differential selection and linkage patterns between the sexes. From the start of Region 2-NP to BP3 (12.2 Mb), there was a distinct difference in zygosity between males and females (Females: 11 HOM ALT, 4 HET, 1 HOM REF; Males: 10 HOM ALT). Therefore, males are out of HWE; based on the female AFs, we would expect 5 HOM ALT, 3 HET and 2 HOM REF in males. The region delineated by BP3 to BP4 overlaps with a high density of SNPs showing signatures indicative of sex-differential selection (12.2 Mb to 13.1 Mb). The only breakpoint consistent between males and females (BP4) also distinguishes the break in LD in males, the drop in gene density, and a break in derived ancestry estimates of 100bp windows (Supplementary Figs. 13, 19, 22). Taken together, this localises our candidate region to the beginning of Region 2-NP to BP4 as both selected in males in a natural population and associated with colour in our breeding design.

## DISCUSSION

Overall, our results reveal a surprising genetic architecture for colour pattern diversity in guppies. Even though the differences in colour pattern between the Iso-Y lines are Y-linked (i.e. inherited faithfully from father to son), consistent genetic differences are most pronounced on an autosome, LG1. Analysis of the phenotype data showed that the breeding design was successful in producing four distinct Iso-Y lines with marked differences in colour pattern. Using these unique Iso-Y lines, we delineated three regions on LG1 that were consistently different among the differently coloured lines, and within these regions we identified many genes known to be linked to colour in model species. Then, using extensive WGS from the source population, we further highlight just one of these regions (Region 2-NP: 11.1 - 15.8 Mb), as having strong linkage and significant local ancestry, finding that a large and variable haplotype is maintained in nature.

It has hitherto been presumed that Y-linked colour pattern genes in the guppy occur on the sex chromosome, LG12. Our results from LG12 do not show consistent differences between the Iso-Y lines. If colour pattern traits are fully Y-linked, and each Iso-Y line’s unique Y haplotype is fully inherited through the patriline, we would expect to see strong differentiation at the terminal region of LG12. All guppy chromosomes are acrocentric and show evidence of male heterochiasmy ^30,58,64^; yet recombination events are particularly rare on LG12, implying that LG12 may decompose at a slower rate compared to autosomes. Indeed, although we identified genetic differentiation on LG12 between the Iso-Y lines, the differentiated regions were not consistent. This suggests that the breeding design may have been unsuccessful in fully breaking down the Y into the constituent parts responsible for colour polymorphism. However, no individual part of LG12 was consistently differentiated among the lines, suggesting a region associated with colour is not on LG12. Using the WGS data, we saw a signature of distinct ancestry at the terminal end of LG12 where we expect the SDL to be located, and another at a region between 5.6 Mb to 7 Mb, part of which has also been previously identified as sex-linked ^31^. Overall, LG12 regions associated with linkage or local ancestry within the natural source population are small, or are associated with the SDL, and notably, there is limited evidence of any obvious colour candidates in these regions.

Our hypothesis is that colour pattern is controlled by the epistatic interaction between Y-specific regions and our candidate region on LG1 in this population. Y-autosome epistasis has been reported previously in other systems. For example, polymorphisms on the gene-poor Y chromosome in *Drosophila* spp. have been shown to differentially affect the expression of hundreds of X-linked and autosomal genes, specifically those that are highly expressed in males and with clear fitness related functions in males (e.g. spermatogenesis and pheromone detection ^65,66^). Here, we found several putative male-specific fitness trait genes on LG1, as well as colour patterning genes. Moreover, recent analytic models predict that Y-autosome epistasis can result in the spread of male beneficial but female deleterious mutations on autosomes, resulting in sexually antagonistic selection (SAS), because of the unique paternal transmission of Y chromosomes (the ‘Father’s curse’ ^67^). We see differences in selection between the sexes on LG1 ‘Region 2-NP’, albeit this region did not appear in our previous analysis exploring consistent differences between the sexes across multiple populations ^31^. Finally, we find that males are not in HWE in the region from 11.1 Mb to 13.1 Mb (start of Region 2-NP to BP4), which could be indicative of sex-biased survival in guppies. Guppy adult sex-ratios, are indeed, generally female-biased ^68–71^, and show a stronger female-bias in upstream low-predation (LP) environments, albeit with sampling variability ^68,71^.

Looking at the phenotype data, we observed that Iso-Y9 was the most differentiated along DF1, but it was also the most variable. At the genome level, Iso-Y9 harboured unique diversity on LG12. Given these results, we can hypothesise that colour pattern distinctiveness is driven by autosomal regions on LG1, but variability is driven by Y-specific regions on LG12. This suggests that guppy males may be able to produce particular novel colour pattern traits via divergence on autosomes, and regulate variability via the Y. It follows that divergence on the Y is potentially able to produce subtle changes to pre-existing autosomal variation in order to produce objectively novel colour patterns.

It has been argued that selection must be extremely strong, or unrealistic, for SA alleles to be maintained on the autosome ^72^. It is also contended that sex differences on autosomes are artefacts caused by duplications or translocations onto the Y chromosome and are thus not due to SAS ^73^. We found no evidence of reduced mapping quality in LG1. Nor did we find evidence that our candidate region on LG1 is misassembled; we previously found strong Hi-C contact across the chromosome, although the genome assembly is derived from an individual from a different population ^31^. We also found no evidence that a translocation from LG1 to LG12 had occurred uniquely in this population in our SV analyses, and did not observe elevated interchromosomal LD (Supplementary Fig. 23). Moreover, linkage along chromosomes, including LG1, has been observed in several different populations and mapping crosses suggesting this is unlikely to be the case ^30,58,74^.

We find no evidence for large structural rearrangements (such as inversions) underlying the maintenance of our candidate region, LG1 ‘Region 2-NP’. We also did not detect any relationship with our predicted conserved recombination points and increased GC or TE content. Therefore, exactly which molecular traits are driving this haplotype structure is unclear. By examining phased heterozygous individuals it is, however, apparent that crossing-over is restricted to key points along the haplotype and overall linkage is maintained in the population. Evidence of increased LD may be indicative of the potential SAS in the LG1 region. Two-locus models have shown that admixture between the sexes with differing allele frequencies can produce LD due to a correlation in the selective forces acting at different loci in the two sexes ^75^. On the other hand, models of balancing selection show that balanced variants produce diversity patterns similar to those caused by positive selection ^76^, and simulations suggest that balancing selection alone can maintain high-LD, and in particular, high divergence between colour phenotypes ^77^. Our data suggest LG1 Region 2 represents a good candidate for autosomal SAS, but confirmation of this hypothesis requires larger sample sizes _78_.

Early research found that the linkage between colour traits and the Y was under selection, with increased Y-linkage for colour traits in downstream, high-predation (HP) environments and X-linkage in the upstream low-predation (LP) environment. Whether Y-linkage varies by predation environment has been indirectly studied across Trinidad, where females treated with testosterone exhibited more colour patterns in LP populations compared to their HP counterparts ^79,80^. Such observations are also consistent with an autosomal genetic component of colour pattern traits that are under weakened selection in LP environments ^81^. Other researchers claim that the size of different aged strata of LG12 differ between HP and LP, with increased Y-linkage in LP ^33^ (see also ^34,35^), but these results have not been repeatable ^30–32^. Our candidate region on LG1 may explain differences in Y-linkage between predation ecotypes. Specifically, we identified a region between 12.2 Mb to 13.1 Mb with reduced diversity in males, compared to females, which also corresponded with a break in male patterns of LD. Previous analysis of molecular convergence between HP and LP populations identified two outlier windows in this area (12.17-12.18 Mb and 12.21-12.2 Mb), indicating that this particular region may be under divergent selection for HP-LP phenotypes ^82^. We further compared WGS data from other populations across Trinidad at LG1, and detected a strong signal of HP-LP association within 12.2 Mb to 13.1 Mb, but the large haplotype structure is unique to the populations studied here (Supplementary Fig. 24). This suggests that this region may be involved in the differential selection of colour phenotypes depending on predation regime.

## Conclusions

Based on our Iso-Y lines and natural population data, we hypothesise a Y-autosome epistatic genetic architecture for guppy colour traits. This architecture may be particularly well-suited to a trait under both SAS and NFDS, such as guppy colour. Models suggest that sex-linked polymorphism can only be maintained by natural selection in unusual genetic systems, where the maintenance of Y-linked variation in Y-autonomous models involves frequency-dependent selection, or interactions with other chromosomes ^83^. Recognition of the importance of epistasis in responding to selection is growing and epistasis may be responsible for an increased/non-linear rate of adaptation in natural populations or artificial selected lines ^84^. Additionally, having loci under NFDS not physically linked to the Y can shield them from increased drift experienced on the Y, allowing for higher levels of variation to be maintained ^28^. Moreover, although SA theory predicts that colour genes should become tightly linked to the SDL, such a situation has actually only been observed in a few teleost species ^24^. Finally, for NFDS to operate, we would hypothesise that there should be a balance between keeping coadapted alleles together, and breaking them apart to create variation. Together, this suggests that Y-autosome epistasis acting on a diverse autosomal haplotype presents a feasible hypothesis for the maintenance of colour pattern traits. The Iso-Y lines offer a unique resource to explore interactions between LG1 and LG12, and also to further distinguish the role of the three different regions on LG1 by performing focussed crosses between Iso-Y lines. Future studies should also aim to fully characterise the diversity of LG1, including additional long-read sequencing of multiple individuals from both LP and HP environments in order to determine the reservoir of variable haplotypes present in this region.

## MATERIALS AND METHODS

### Generation of the Iso-Y lines

The Iso-Y lines were kindly provided by AE Houde, who established them by choosing male lineages in which colour pattern on the body was strongly Y-linked. Each line was founded by a single male drawn from the ‘Houde’ tributary of the Paria River in Trinidad (Trinidad National Grid System: PS 896886). Males from this tributary are known to show strong Y-linkage ^19^. Each Iso-Y line was maintained by breeding males with colour patterns similar to that of each line’s founder; hence, the colour patterns of the Iso-Y lines are ecologically relevant. Every generation, males were mated to females sired by males from the same line, or, every 2-3 generations, backcrossed into the stock population. The lines have been maintained at Florida State University since 2012.

### Colour pattern phenotype analysis

#### Photography

We used multispectral digital photography (Sony A7 with full-spectrum conversion; Nikon 80mm f/5.6 El-Nikkor Enlarging Lens) to capture human-visible and ultraviolet images of males from each Iso-Y line (Iso-Y6 _n_ = 41; Iso-Y8 _n_= 48; Iso-Y9 _n_= 42; Iso-Y10 _n_= 42). Fish were lightly anesthetised by immersion in a Tricaine mesylate (Pentair) solution and placed on a clear petri dish above a grey background with the left side of the body facing upwards. A soft tip miniature paint brush was used to raise the dorsal fin, flare the caudal fin, and lower the gonopodium so that these appendages were visible. Fish were illuminated by four metal halide lights (Hamilton, 6500 K bulbs) that simulate the natural photic environment. A size standard and two full-spectrum colour standards (grey – 20% reflectance, white – 99% reflectance; Labsphere) were included beside the fish. Glare and shadow were minimised by placing a diffuser (cylinder of 0.015” polytetrafluoroethylene) around the fish and colour standards. We photographed each fish once in the human visible spectrum (Baader UV/IR cut / L-Filter) and once in the UV spectrum (Baader U-Venus-Filter 350nm).

#### Morphometrics

Morphometrics were performed using the *TPS Series* software ^85^. We used *tpsDig2* v2.31 ^86^ to set the image scale using the size standard, and to place landmarks around the perimeter of the fish. Following Valvo et al. (2020) ^40^, we placed seven ‘traditional landmarks’ at the tip of the snout, the anterior dorsal fin attachment, the posterior dorsal fin attachment, the dorsal caudal fin attachment, the ventral caudal fin attachment, the posterior gonopodium attachment, and the anterior gonopodium attachment. We then placed 55 ‘semi-landmarks’ at approximately even intervals between the traditional landmarks. We used sliding of semi-landmarks to minimise any shape variation resulting from unequal distribution of semi-landmark placement. Next, *tpsSuper* v2.05 ^87^ was used to generate a ‘consensus shape’ representing the average shape of the males in all 346 photos (173 males * 2). Images of each individual were ‘unwarped’ to this consensus shape, thereby mapping every pixel from the original image to an analogous location on the consensus shape.

#### Colour Measurement

Analysis using the Colormesh pipeline ^40^ was performed in R v4.0.2. We first performed Delaunay triangulation, which is used to reconstruct a complex shape (i.e. the shape of the unwarped fish) using a concise number of points distributed across the surface of that shape. We then measured the average colour of pixels in a radius around each of these sample points. We used cross-validation to determine the optimal number of Delaunay triangulations (more triangulations result in more granular colour pattern data) and sample circle radius (see below). At each sample point, we extracted linear colour measurements for four colour channels: R (red), G (green), and B (blue) and UV. We accounted for any minor fluctuations in the lighting environment by calibrating the colour values for each channel by subtracting the average deviation of colour measured in the photo on the white and grey colour standards from the known reflectance values of those standards.

#### Colour pattern differences among Iso-Y lines

We used Discriminant Analysis of Principal Components (DAPC) to describe the properties of colour pattern as this analysis is recommended for characterising the properties of groups using high dimensional data sets. We used the *dapc* function in *adegenet* v2.1.3 ^41^ to perform a Principal Components Analysis (PCA), followed by a discriminant analysis to define the linear combinations of PC scores that minimise within and maximise between group variances.

We used DAPC in a cross-validation framework to determine the scheme for capturing colour data that allowed us to best discriminate among the Iso-Y lines ^40^. We used cross-validation to determine the optimal number of PC’s to retain, number of Delaunay triangulations to perform, and sample circle radius. Using the *xvalDapc* function, we performed DAPC on training and validation data, and examined the average proportion of successful assignments for a varying number of retained PC’s. We defined the training and validation populations each as 50% of the individuals from each line. We set the maximum number of PC’s to retain for cross-validation as *n*.*pca* = 173 (the number of fish), with cross-validations performed at 17 different retention levels in increments of 10 PC’s (up to 170). We performed 100 replicates per PC retention level. The average proportion of successful placements was maximised (at 98.8% successful) by retaining 10 PC’s, using four Delaunay triangulations, and a sample circle radius of one pixel. Consequently, we used this sampling scheme to measure colour pattern for all phenotypic analyses, and retained 10 PC’s for DAPC. This scheme resulted in 9904 colour measurements per fish (four colour channels * 2476 sampling locations), with colour averaged over 5 pixels per sample point.

To visualise the colour variation summarised by the discriminant functions, we generated heat maps depicting the correlations between position on each discriminant function and colour for each colour channel at each sampling location.

#### Comparing colour pattern differences among Iso-Y lines

To determine whether the Iso-Y lines have robust phenotypic differences, we used a permutational MANOVA to compare mean colour measures among lines. We first reduced the dimensionality of the colour pattern data for this analysis using PCA. We retained the first 17 PC’s to summarise the greatest amount of variation in the data while keeping the ratio of observations to variables greater than 10:1 for our subsequent multivariate analyses. These PC’s together explained 59.3% of the total variation in colour measurements. We then compared PC scores among lines using the *Adonis* function in *vegan* v2.5-6 ^88^, which computes the pairwise distances among observations and then performs a permutation test (randomly assigning labels among factors) to partition the distance matrix among sources of variation. P-values are then calculated using pseudo-F ratio tests. We used Euclidean distance and created 10 000 permuted samples per test. We additionally visualised the differences in phenotype among Iso-Y lines by generating density plots of each Iso-Y lines for all 17 PC’s used in permutational MANOVA (Supplementary Fig. 2).

#### Comparing colour pattern variance among Iso-Y lines

To quantify total within-line variance in colour pattern, we calculated the trace of the variance-covariance matrix among fish within a given line (across all colour channels and sampling locations). We then used permutational t-tests to determine whether phenotypic variance differed between each pair of lines, using the *sample* function. We permuted the line labels for each whole-fish colour pattern, thereby accounting for the fact that different colour measures on the same fish are not independent of one another. We created 10 000 permuted samples per test, and performed two-tailed tests.

### Genomic library preparation methods and variant genotyping

#### Pool-seq of the Iso-Y lines

Genomic DNA was extracted from 48 males of each Iso-Y line using an ammonium acetate extraction method from caudal peduncle ^89^. DNA quality was assessed using gel electrophoresis and DNA concentration calculated using Quant-iT Picogreen dsDNA reagent (Invitrogen) on a Glomax Explorer Microplate reader (Promega). DNA of each individual was diluted to 18-22ng/µl and 4µl of each added to a pooled sample per Iso-Y line. Pools were cleaned using a NEB Monarch clean up kit, with final concentration and quality of each confirmed using a Nanodrop ND-1000 (Thermo Fisher) and a Bioanalyser (Agilent). Library preparation and sequencing was performed at the University of Exeter Sequencing Service, using a NextFlex RAPID PCR-free library preparation protocol. Each of the libraries were sequenced across multiple lanes on a HiSeq 2500 in standard mode, with a 125bp paired-end metric (Supplementary Table 12).

Raw reads were cleaned using cutadapt v1.13 ^90^. Reads were aligned to the guppy genome ^31^ with bwa mem ^91^ and converted to sorted bam alignment format with samtools v1.9 ^92^. Coverage was calculated using qualimap v2.2.1 ^93^. Variants were called using Freebayes v1.3.1 ^94^ with GNU parallel ^95^ by chunking the alignment files into regions based on coverage using sambamba v0.7 ^96^. Freebayes was run with the options: *--pooled --use-best-n-alleles 4 -g 1000*. The raw variant output was filtered for bi-allelic SNP variants at QUAL > 30 and DP > 10, followed by a max-missing filter of 80% applied to each pool separately followed by the application of a minor allele frequency (maf) of 25% applied across all pools using vcftools ^97^. The VCF file was used as input to poolfstat ^98^, where the variants were filtered to exclude sites at <30x minimum coverage and <500x maximum coverage per pool, with a minimum read count per allele of 10 (Final SNP set = 3,995,905 SNPs).

#### Whole genome sequencing (WGS) of wild-caught individuals

For the WGS of 26 wild-caught guppies, we sequenced individuals from the Paria river (n=9) and used these with previously available data ^31^ (Supplementary Table 13). Genomic DNA was extracted using the Qiagen DNeasy Blood and Tissue kit (QIAGEN, Hilden). DNA concentrations ≥ 35ng/μl were normalised to 500ng in 50μl and were prepared as Low Input Transposase Enabled (LITE) DNA libraries at The Earlham Institute, Norwich. LITE libraries were sequenced on an Illumina HiSeq4000 with a 150bp paired-end metric and a target insert size of 300bp, and were pooled across several lanes so as to avoid technical bias with a sequencing coverage target of ≥10x per sample. Data from the LP Marianne river was previously generated (n=17) ^31^. Although the Paria and LP Marianne guppies are sampled from different sites, there is strong evidence that gene flow occurs between the populations occupying the upper reaches of these rivers ^99,100^. We found that genome-wide mean *F*_ST_ calculated between populations in this dataset was 0. Importantly, there was no effect of river on haplotype structure (Supplementary Table 13). Data processing followed previous methods using the GATK4 Best Practices pipeline (https://github.com/josieparis/gatk-snp-calling). Final filtering of the VCF file included filtering for bi-allelic SNPs at a minimum depth of 5 and a maximum depth of 200, removing 50% missing data and application of a 10% maf filter (Final SNP set = 1,021,495). Variants were phased individually with Beagle v5.0 ^101^, which performs imputation and phasing, and then phased again using Shapeit v2.r904 ^102,103^ making use of phase-informative reads (PIR) ^104^.

#### Long-read sequencing

To assist with the phasing of variants called from short-read data and to detect structural variants (SVs) we generated 20Gb of long-read Pacbio data from one of the Iso-Y individuals (Iso-Y6). High molecular weight DNA was extracted using DNeasy Blood & Tissue Kit (Qiagen, The Netherlands) with modifications (10x Genomics Sample Preparation Demonstrated Protocol and MagAttract HMW DNA Kit handbook). Data were sequenced on 3 SMRT cells of a PacBio Sequel at the University of Exeter Sequencing Service. For phasing, reads were aligned to the guppy genome using minimap2 ^105^. Genotypes were phased using whatshap v0.18 ^106^ using the reference genome and the long-reads to lift phasing information.

### Delineation of Iso-Y-specific blocks and analysis of differentiated regions

Pairwise *F*_ST_ was calculated using the Anova *F*_ST_ method implemented in poolfstat ^98^. Per SNP *F*_ST_ values were compared pairwise between each Iso-Y line. To identify chromosomes with the highest mean variance in *F*_ST_ differentiation across our dataset, mean Z-scores of each PC were summarised for each chromosome ^43^. These were calculated using the *prcomp* function in R, centering and scaling the results.

To delineate line-specific blocks apparent from the distinct patterns of differentiation observed in the *F*_ST_ analysis, we evaluated the allele frequencies (AFs) of each line. AFs were extracted from the Pool-seq object generated by poolfstat and were polarised to Iso-Y9 (the line which showed the highest genetic differentiation). To explore delineation breakpoints in the AFs, we adopted the use of change point detection (CPD) analysis. CPD was conducted in R using *changepoint*^107^. We used individual AFs from each line as input to detect mean changes, using the BinSeg method and SIC criterion. The number of changepoints identified is Q; in cases where several change points were detected, we increased the value of Q to 10. To add support to the change points detected using the AFs, and to ensure we were not missing any additional breakpoints, we also used PC1 *F*_ST_ scores as input. In cases where multiple change points were detected within close vicinity of one another, caution was taken to delimit the smallest region in each case; this was to correctly identify the minimum unit of inheritance. Within the identified CPD regions, we further inspected the segregation of Iso-Y line alleles responsible for driving differentiation by plotting the AFs of each identified region.

To assess diversity among the Iso-Y lines, π was calculated from the Iso-Y line allele frequencies on a per base pair basis ^108^. To summarise the among-Iso-Y line variation, per SNP π values were used as input to PCA, calculated in R using the *prcomp* function.

For functional gene annotation, we extracted the regions of interest from the guppy genome (ENA: GCA_904066995) using samtools faidx ^92^ and aligned them to the previous guppy genome assembly (Ensembl: GCA_000633615.2) using minimap2 ^105^ and pulled the uniprot gene IDs from the annotation using biomaRt ^109^.

We used the curated liftover for LG12 for all analyses and plots of LG12 ^110^.

### Exploration of Iso-Y differentiated regions in natural populations

WGS data comprising 26 wild-caught individuals was used to explore the identified Iso-Y regions in natural guppy populations. Intrachromosomal linkage disequilibrium (LD) was calculated among polymorphic SNPs for LG1 and LG12. Invariant positions were removed using bcftools ^111^, variants were thinned at 5kb intervals and *r2* values were calculated in plink v1.8 ^112^; bwh.harvard.edu/plink), outputting a square matrix for plotting using LDheatmap ^113^.

Shifts in localised heterogeneity have been explained as a potential artefact of chromosomal inversions or long maintained haplotypes. We analysed patterns arising from changes in local ancestry in the natural data using *lostruct* v0.9 ^56^. Local PCAs were run with three PCs mapped onto three MDS. Mapped eigenvalues > 0.01 were assessed for saturation before defining significant MDS. Significant outlier windows of each MDS were defined by first calculating 3 standard deviations of the mean of the MDS distribution after trimming the 5% tails of each distribution, followed by returning all windows at the extremes of the distribution. To balance loss of power and sensitivity, local PCAs were assessed in both 10 bp and 100bp windows (Supplementary Fig. 22). We considered the start and end positions of the signature of each MDS as the last region where we found 3 or more adjacent neighbouring windows.

Population genetics statistics were calculated along LG1 and LG12 using PopGenome v2.7.5 ^114^. Intersex-*F*_ST_, D_xy_ and π were calculated in non-overlapping 1kb windows. Intersex *D*_a_ was calculated as D_xy_ - female π. Outliers were considered if the regions were outside of the upper and lower 95% quantiles of each calculated statistic. GC and repeat content were calculated from the male guppy genome ^31^, and were computed in 1kb windows.

### Structural variants and coverage

For the medium-coverage WGS data derived from wild-caught individuals we used BreakDancer v1.4.5 ^115^ and Lumpy-based smoove 0.2.5 ^116^ using SVtyper ^117^. We also applied these two short-read methods to the Iso-Y Pool-seq data, in addition to Manta v1.6.0 ^118^ as the data were high-coverage and PCR-free. SV outputs were filtered depending on the software: breakdancer SVs were filtered at a confidence score >=99, read group (RG) support >=median RG; manta SVs were filtered at a quality score >=999 and only events where both paired-read and split-read support at values realistic of the overall read depth were retained; for Lumpy/smoove SVs, events that had evidence of both split read and paired end support at realistic read depth values were considered. We excluded SVs <= 10kb. We also made use of our long-read sequencing data from a Iso-Y line individual (Iso-Y6) using sniffles v.1.0.12 ^119^ and PBSV v 2.3.0 (https://github.com/PacificBiosciences/pbsv). Any suspected SVs were manually inspected in the Integrated Genomics Viewer (IGV v2.4.8; ^120^).

Coverage was estimated from bam files using the bamCoverage option of deeptools v3.3.1 ^121^. The first and last 100 kb of each chromosome were trimmed before estimating coverage with a binSize = 50, smoothLength = 75, and an effectiveGenomeSizeof 528,351,853 bp (genome size minus masked and trimmed regions). The ends of chromosomes were trimmed because they often showed high peaks of coverage, thereby distorting the normalised measures across the chromosome. Coverage estimates were normalised using RPGC (reads per genomic context). Bins with normalised coverage > 4 (four times the expected median of 1) were filtered. For the natural WGS data, coverage was averaged within populations. Outputs were converted to bed files with window sizes of 1 and 10 kb by taking weighted means.

### Analyses of multiple haplotypes on LG1

Genotype and haplotype plots were created using a custom genotype plot function in R (https://github.com/JimWhiting91/genotype_plot). For the haplotype-based analysis, we polarised the phased haplotypes to Iso-Y9 (also available https://github.com/JimWhiting91/genotype_plot). Breakpoints between the phases were quantified by identifying the location (in bp) where phase0 switched to phase1, and vice versa. Switchpoints were quantified in males and females and a switchpoint was considered to be conserved when the locations (within 50kb) overlapped in two or more individuals.

Interchromosomal linkage was calculated as above for intrachromosomal linkage calculating *r2* values between LG1 and LG12 using the *--inter-chr* flag in plink, outputting a normal pairwise matrix. Results were plotted in R using custom scripts (https://github.com/josieparis/interchromLD).

## DATA AVAILABILITY

DNA sequencing data of the Iso-Y lines are available at the European Nucleotide Archive (ENA) under the Project Accession PRJEB36506. Whole-genome sequencing data for individuals available on ENA (Project Accession PRJEB10680 and Project Accession XXX). Long-read pacbio data for Iso-Y6 is accessible under Project Accession XXX. All scripts and other data associated with analysis will be made available in an archived github repository (Zenodo, DOI: XXX).

## ACKNOWLEDGMENTS

We wish to thank Anne Houde for the initial collection of the Iso-Y line fish from the Paria River, Jenn Valvo for assistance in the maintenance of the Iso-Y lines, and Sally Lepzinski for helping photograph fish. Thanks to Deborah Charlesworth for experimental insight and suggestions. Computational infrastructure support was provided by The University of Exeter’s High Performance Computing (HPC) facility (ISCA). DNA sequencing was performed by University of Exeter Sequencing Service (ESS). The project was funded by the Natural Environment Research Council (NERC, NE/P013074/1) (JRP, BAF), EU Research Council grant (GuppyCon 758382) (BAF, JRW, MvdV) and the National Science Foundation of the United States (NSF) ISO-1354775 and DEB-1740466 (KAH, MJD).

## AUTHOR CONTRIBUTIONS

JRP carried out molecular work for the WGS data, performed genomic and statistical analysis, interpretation, and wrote the manuscript. JRW assisted with analysis and interpretation throughout. MJD conducted the breeding design, performed phenotyping analysis and wrote parts of the manuscript. JFO assisted with analysis, interpretation and figure preparation. PJP performed the molecular lab work for the Pool-seq data. MvdZ assisted with molecular work and analysis. CWW assisted with analysis of the Pool-seq data. KAH conceived the project, oversaw the breeding experiments, provided analysis and interpretation suggestions throughout. BAF conceived and supervised the project, provided assistance with analysis and co-wrote the manuscript. All authors provided comments on earlier drafts.

## COMPETING INTERESTS

The authors declare no competing interests.

## Notes

### Competing Interest Statement

The authors have declared no competing interest.

