## Supplementary Figures for "A large and diverse autosomal haplotype is associated with sex-linked colour polymorphism in the guppy"

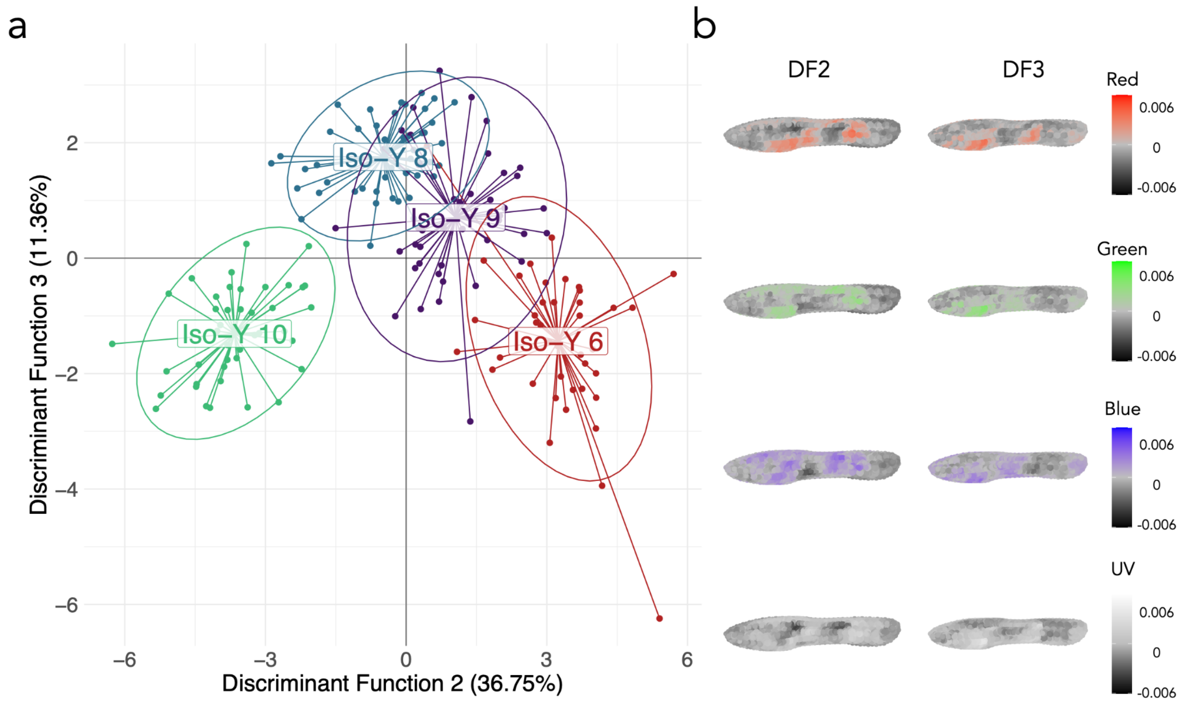


**Supplementary Figure 1**. Discriminant analysis of principal components (DAPC). (a) Scatterplot of discriminant functions 2 & 3. (b) Heatmaps for each colour channel depicting the correlations between colour and at each sampling location and discriminant function 2 or 3.

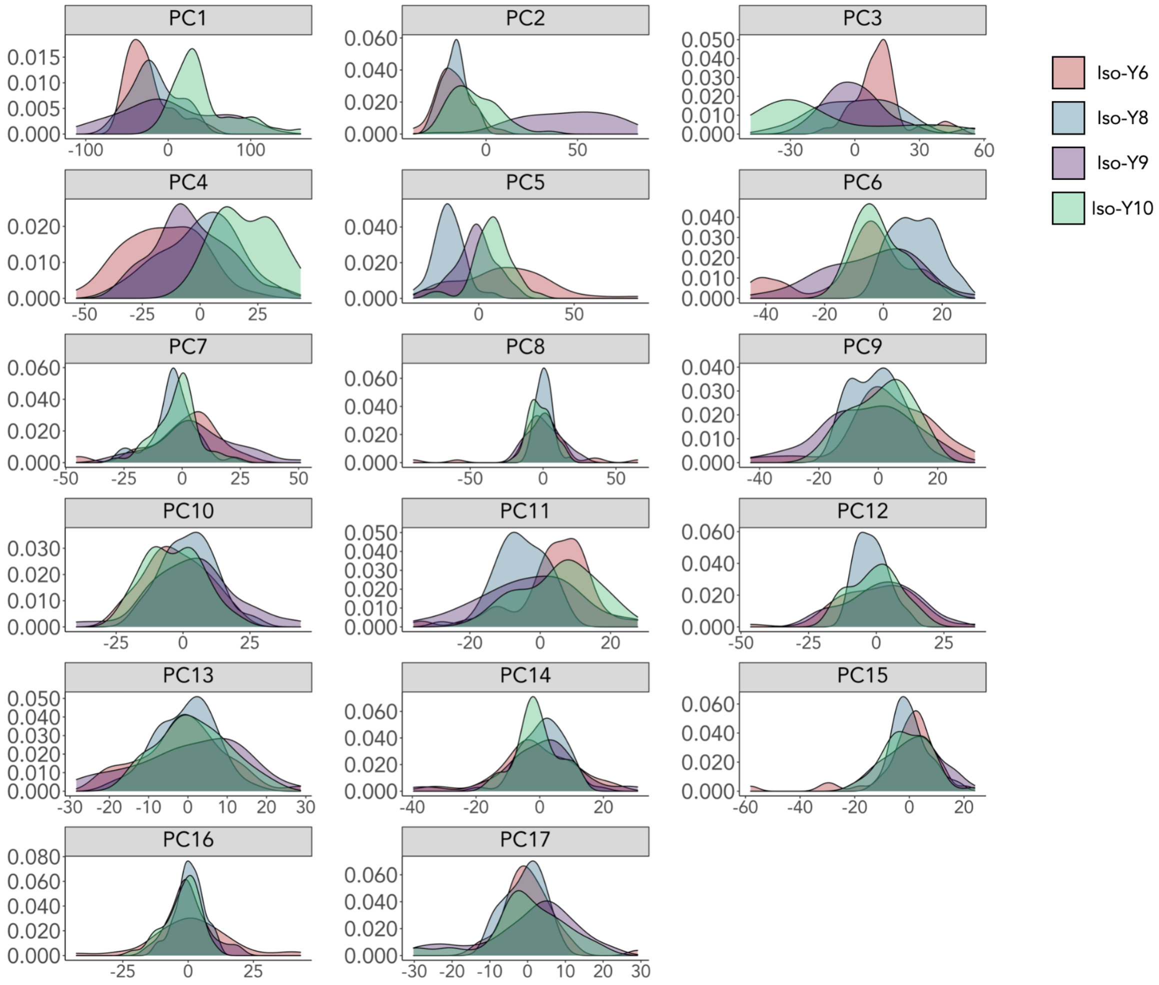


**Supplementary Figure 2.** Density plots of the phenotype of each Iso-Y line for the first 17 principal components.

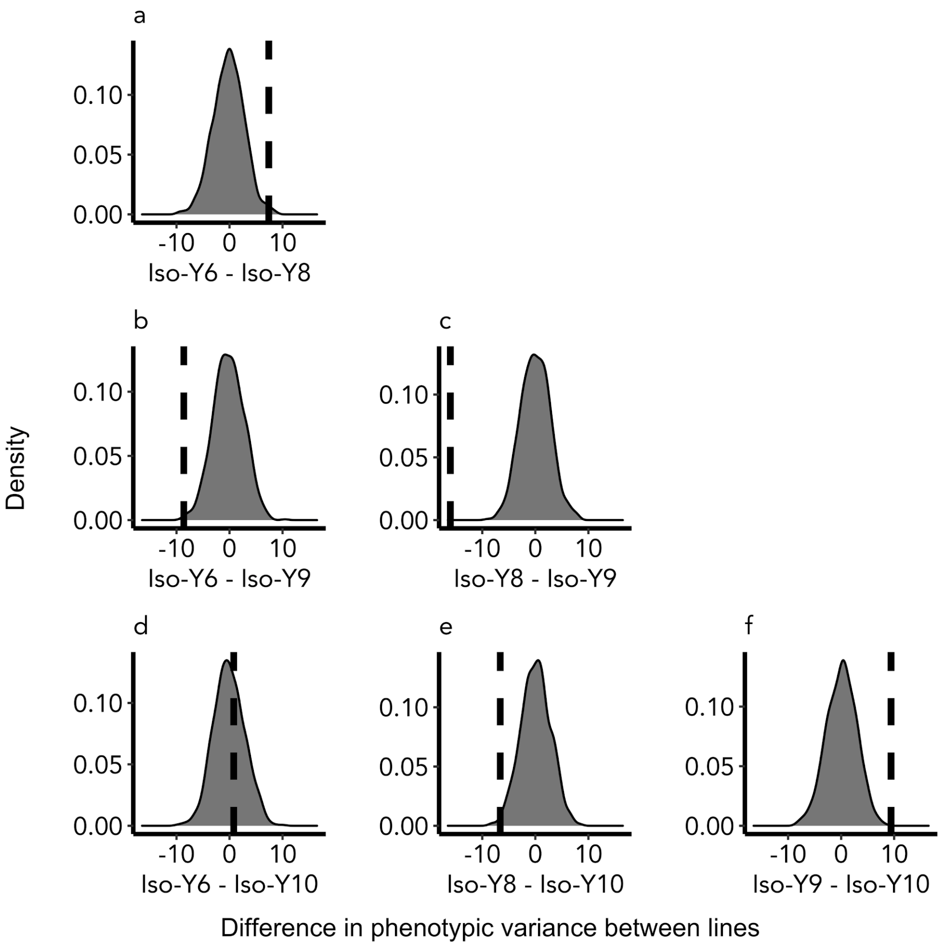


**Supplementary Figure 3.** Observed and permuted differences in phenotypic variance between each pair of Iso-Y lines. The density curve describes the null distribution of variances; the dashed line denotes the observed difference in variance.

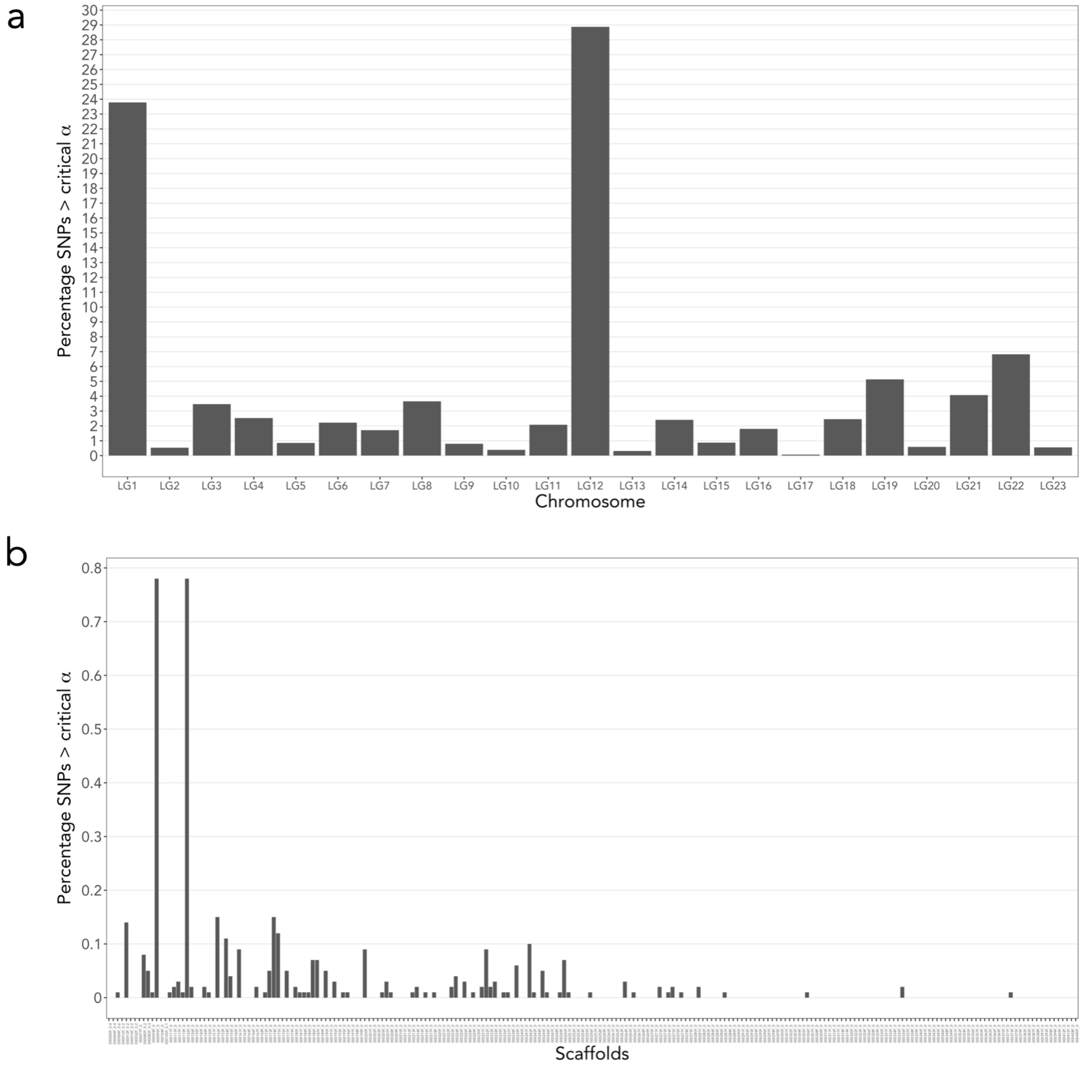


**Supplementary Figure 4**. Percentage of SNPs with a high Z-$F_{ST}$ PC1 above a critical Z-score (Z-$F_{ST}$ PC1 > 3, α=0.05) across the genome: a) per chromosome; b) per scaffold.

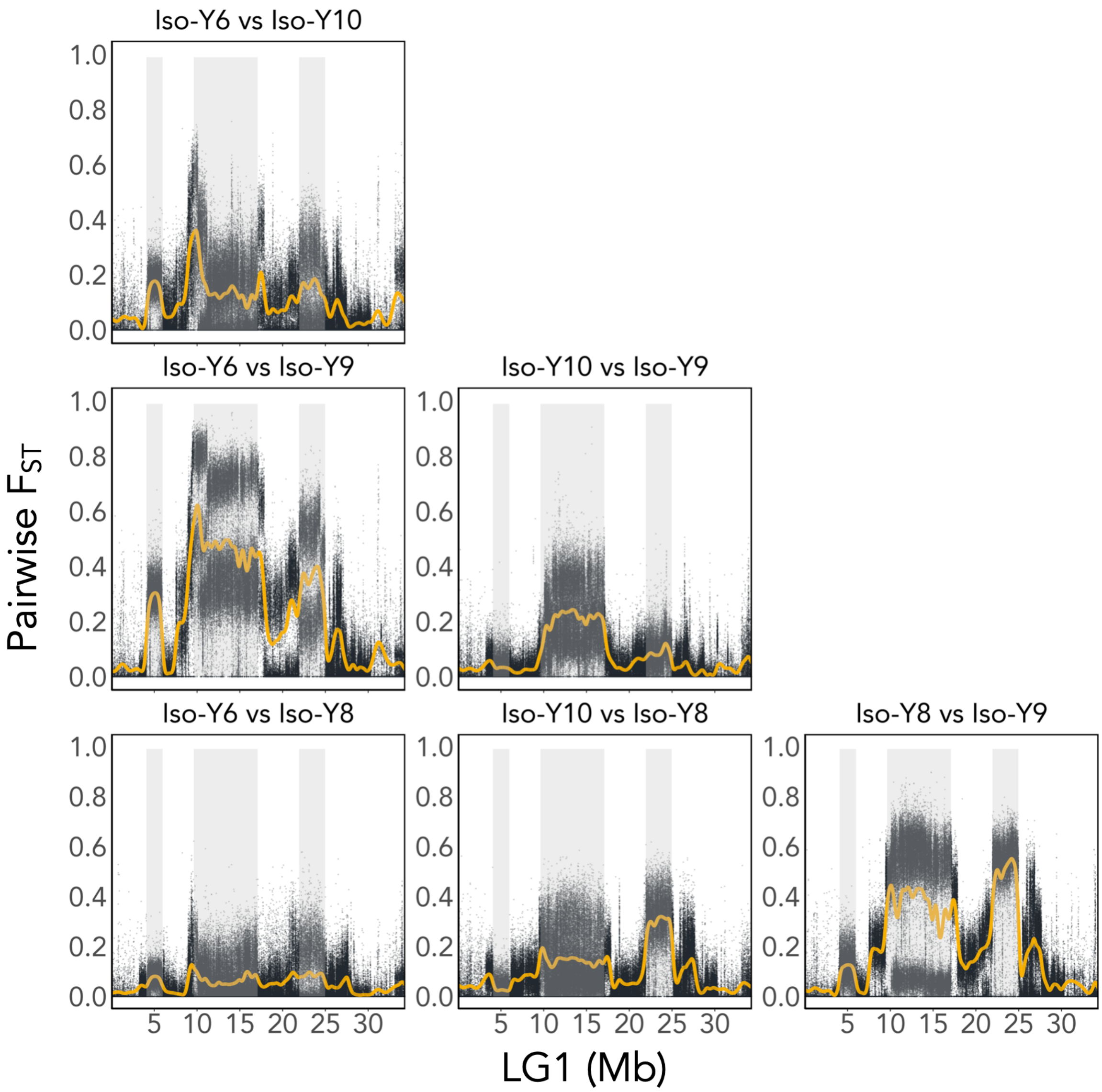


**Supplementary Figure 5**. LG1 Pairwise $F_{ST}$ calculated between the four Iso-Y lines: Iso-Y6, Iso-Y8, Iso-Y9 and Iso-Y10. Yellow lines represent a smoothed spline of the data. Shaded areas represent the three regions identified by change point detection.

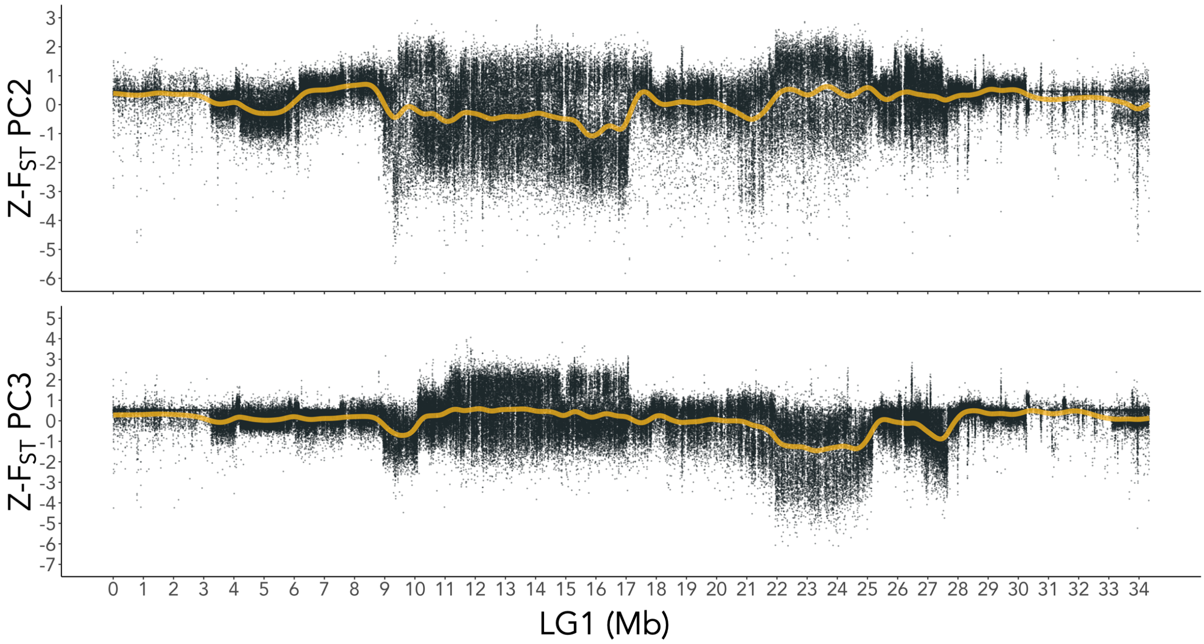


**Supplementary Figure 6**. *Z*-$F_{ST}$ PC2 for LG1; yellow line represents a smoothed spline of the data. PC2 accounted for 17% of the total variance. PC3 accounted for 16% of the total variance.

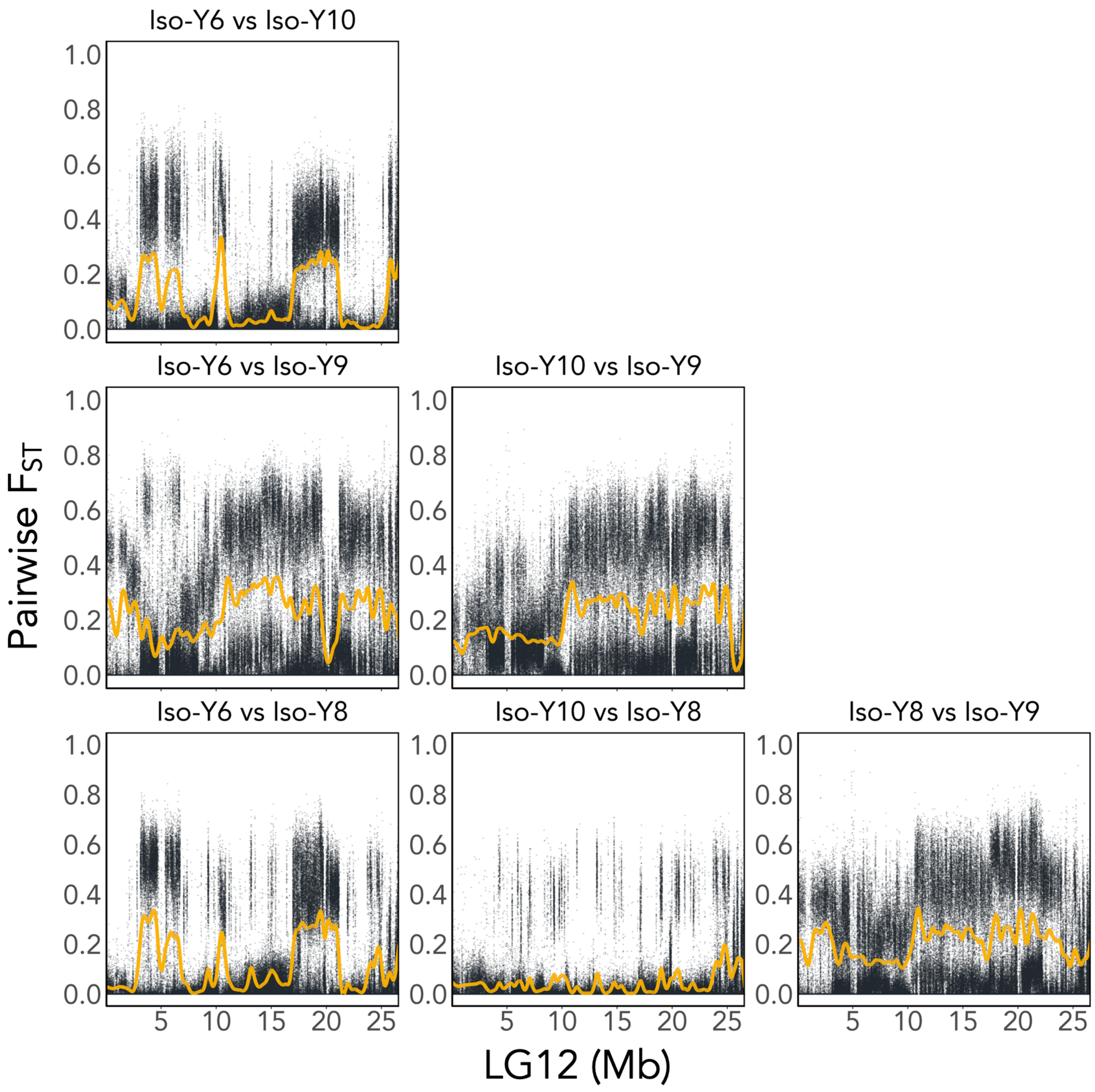


**Supplementary Figure 7.** LG12 Pairwise $F_{ST}$ calculated between the four Iso-Y lines: Iso-Y6, Iso-Y8, Iso-Y9 and Iso-Y10. Yellow lines represent a smoothed spline of the data. No regions were identified by change point detection.

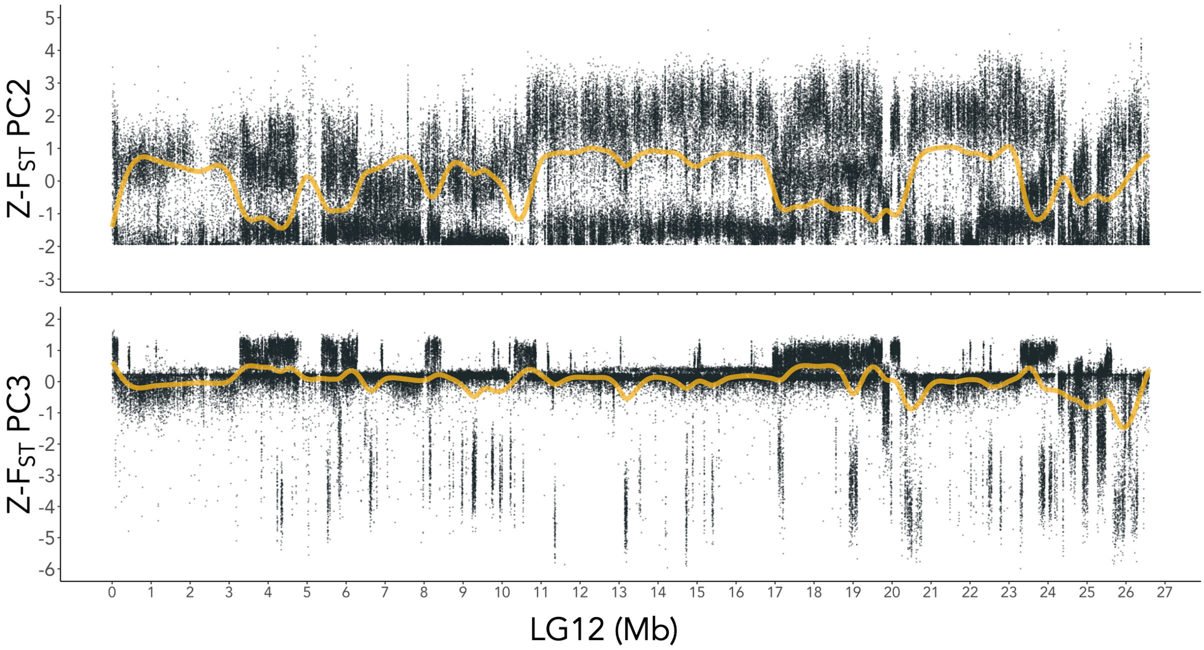


**Supplementary Figure 8.** Z-$F_{ST}$ PC2 and Z-$F_{ST}$ PC3 for LG12; yellow line represents a smoothed spline of the data. PC2 accounted for 30% of the total variance. PC3 accounted for 16% of the total variance.

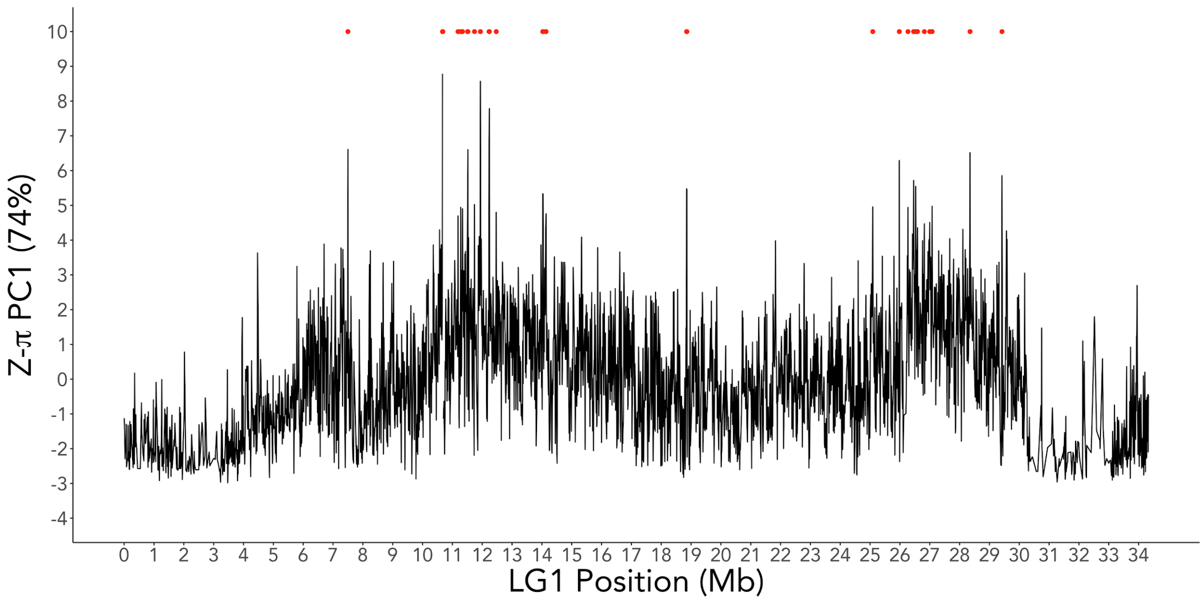


**Supplementary Figure 9.** Z-π-LG1 PC1 (74% of variance) along LG1, which represents the general diversity landscape of the chromosome shared by all Iso-Y lines (SI Table 6). Red points mark 10kb windows with Z-π-LG1 PC1 above a critical Z-score (Z-π PC1 > 4.3, α=0.01).

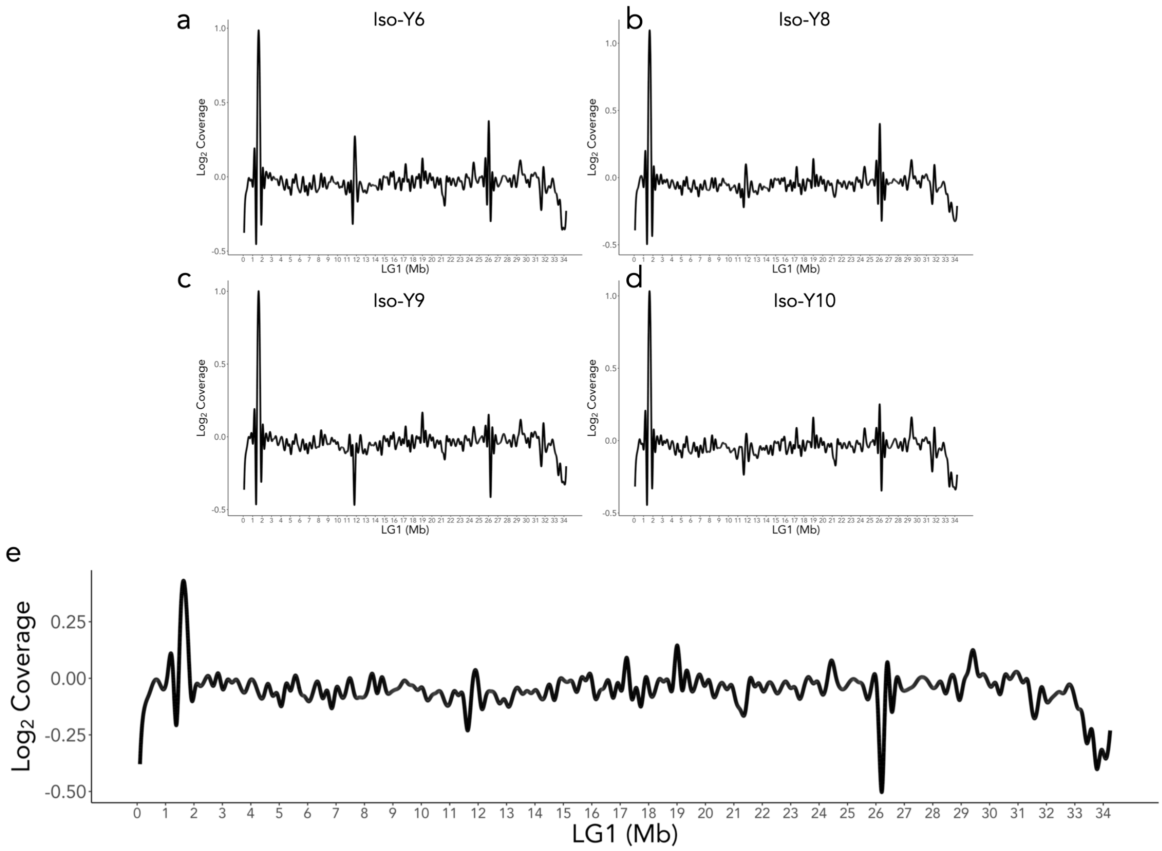


**Supplementary Figure 10.** Coverage calculated across LG1 for each of the four Iso-Y lines: a) Iso-Y6; b) Iso-Y8; c) Iso-Y9; d) Iso-Y10; e) averaged over the mean of all Iso-Y lines.

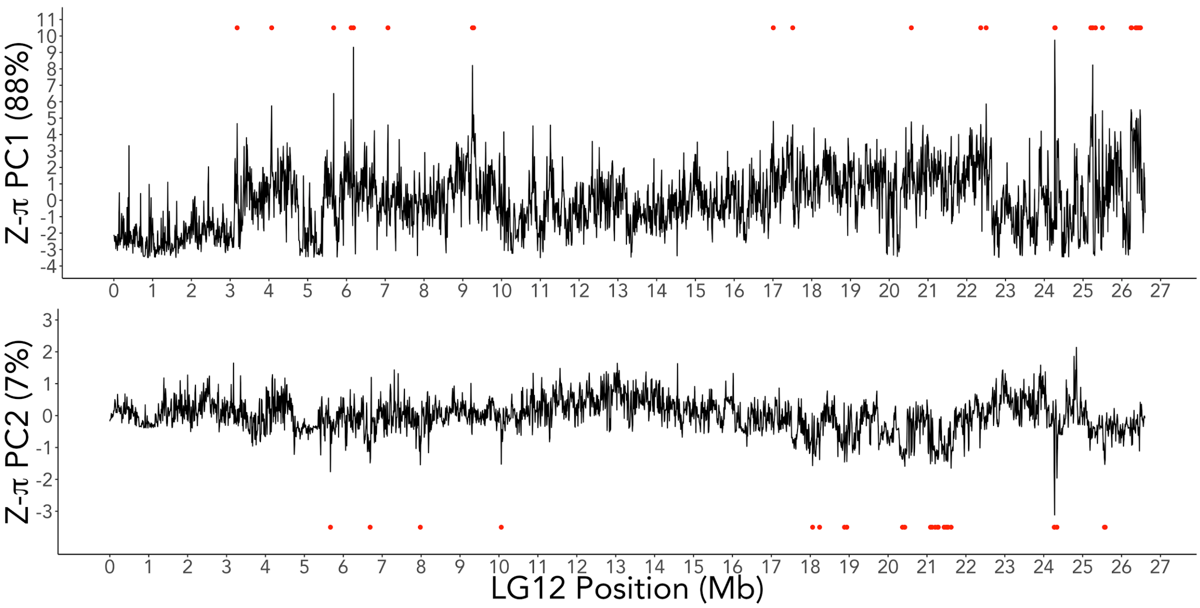


**Supplementary Figure 11.** Z-π PC1 (88% of variance) and Z-π PC2 (7% of variance) along LG12. PC1 represents the general diversity landscape of the chromosome shared by all Iso-Y lines (SI Table 7). Red points mark 10kb windows with Z-π-LG1 PC1 and Z-π-LG1 PC2 above a critical Z-score (Z-π PC1 > 4.6; Z-π PC2 < -1.4, α=0.01). The highest scoring critical window on PC1 was found at 24.27 Mb. Iso-Y9 showed residual variance associated with PC2, where the highest scoring critical window on PC2 was found at 24.28 Mb.

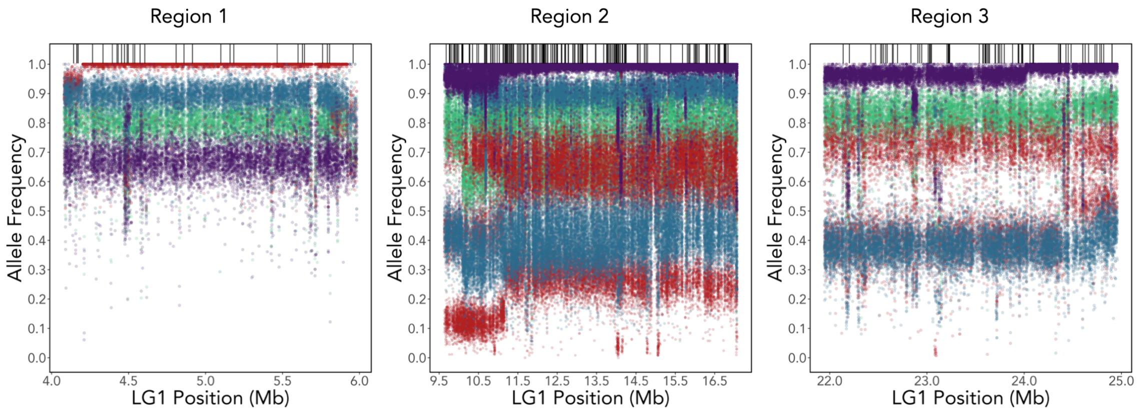


**Supplementary Figure 12.** LG1 polarised allele frequencies for Iso-Y6 (red), Iso-Y8 (blue), Iso-Y9 (purple) and Iso-Y10 (green) for the three identified regions of differentiation: a) Region 1, fixed in Iso-Y6 (coordinates: 4,079,988 - 5,984,584 bp); b) Region 2 (coordinates: 9,627,619 - 17,074,870 bp), fixed in Iso-Y9 and c) Region 3 (coordinates: 21,944,840 - 24,959,750 bp), fixed in Iso-Y9. Further detail of Region 2 and Region 3 can be found in SI Figure 13 & 14.

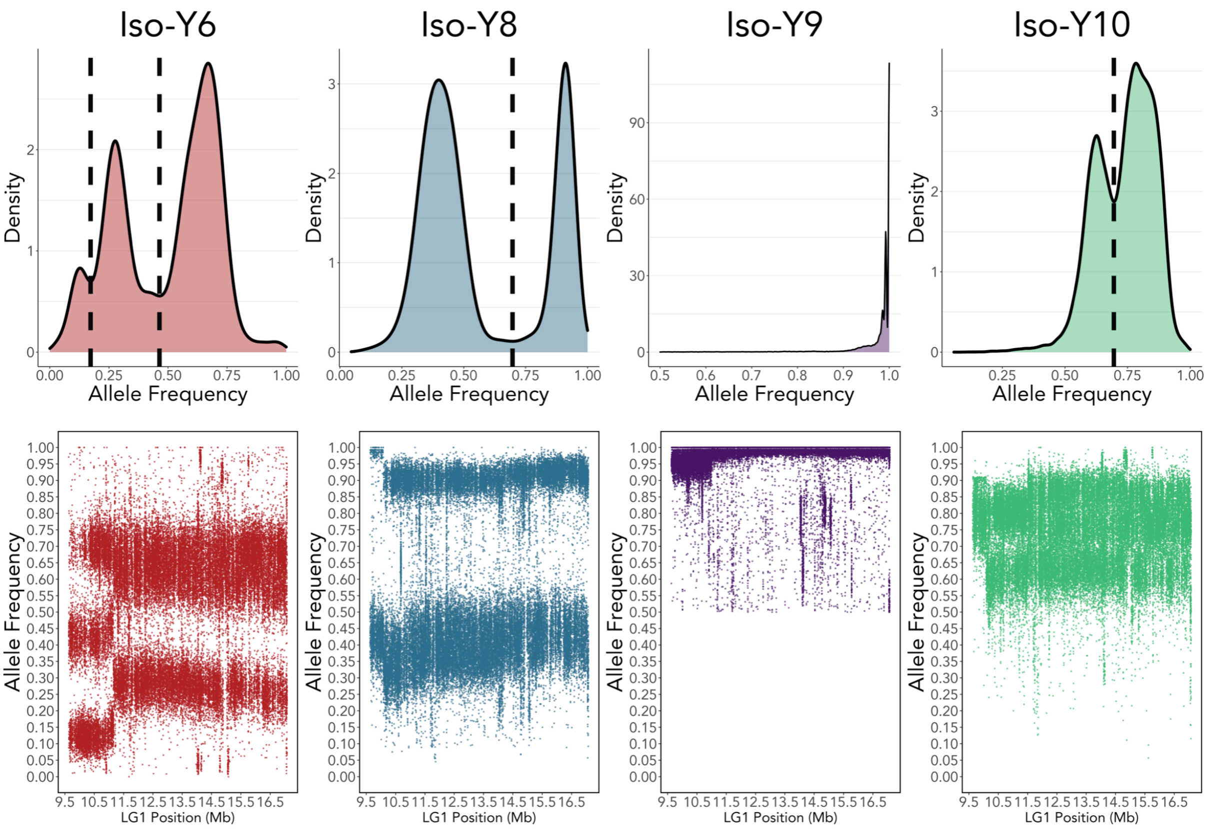


**Supplementary Figure 13.** Allele frequency (AF) density distributions for LG1 Region 2 (coordinates: 9,627,619 - 17,074,870 bp). Iso-Y6 (purple) shows a trimodal distribution of AFs, with three distinct bands of segregating AFs in the first part of the region, and two distinct AF bands for the remainder of the region. Iso-Y8 (blue) and Iso-Y10 (green) both show bimodal AF distribution with two distinct bands of segregating AFs. Iso-Y9 (yellow) shows fixation of the AFs.

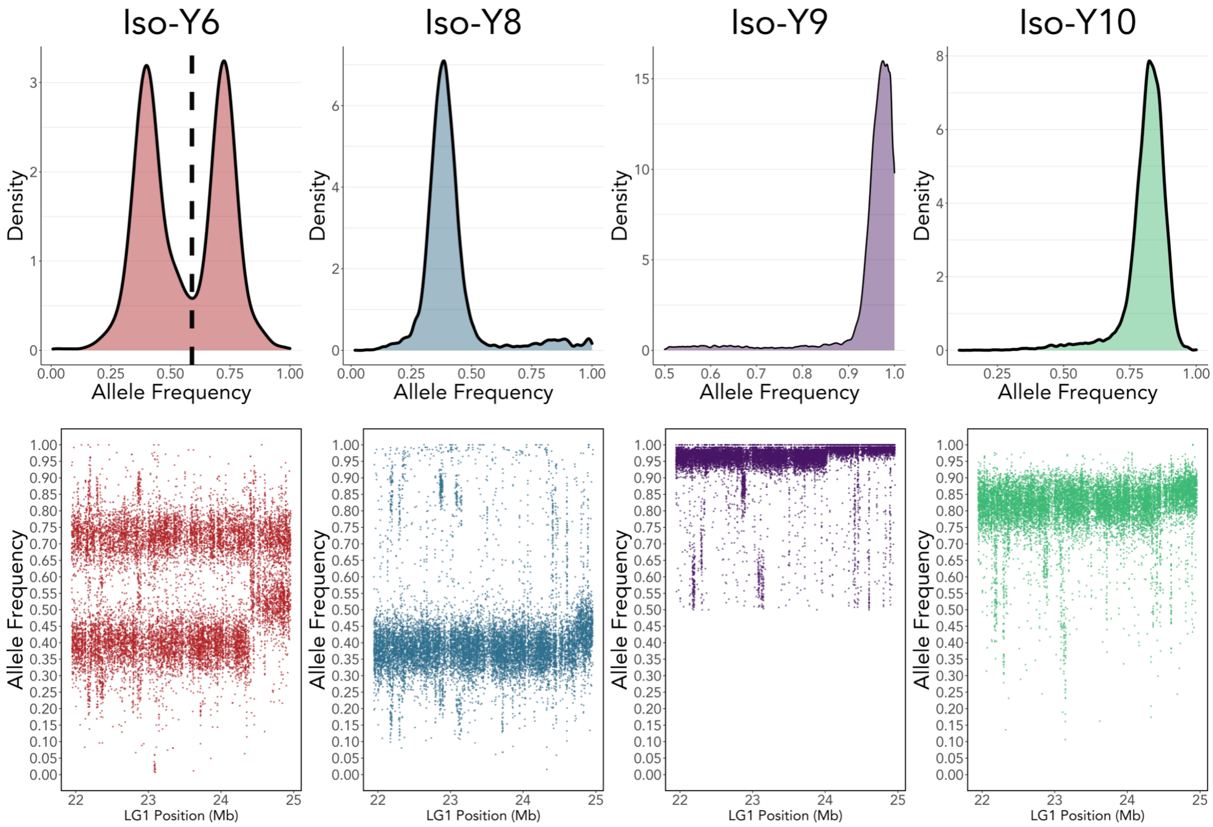


**Supplementary Figure 14.** Allele frequency (AF) density distributions for LG1 region 3 (coordinates: 21,944,840 - 24,959,750 bp). Iso-Y6 (purple) shows a bimodal distribution of AFs, with two distinct bands of segregating AFs. Iso-Y8 (blue), Iso-Y9 (yellow) and Iso-Y10 (green) both show a single AF distribution with one band of segregating AFs. Iso-Y9 (yellow) shows fixation of the AFs.

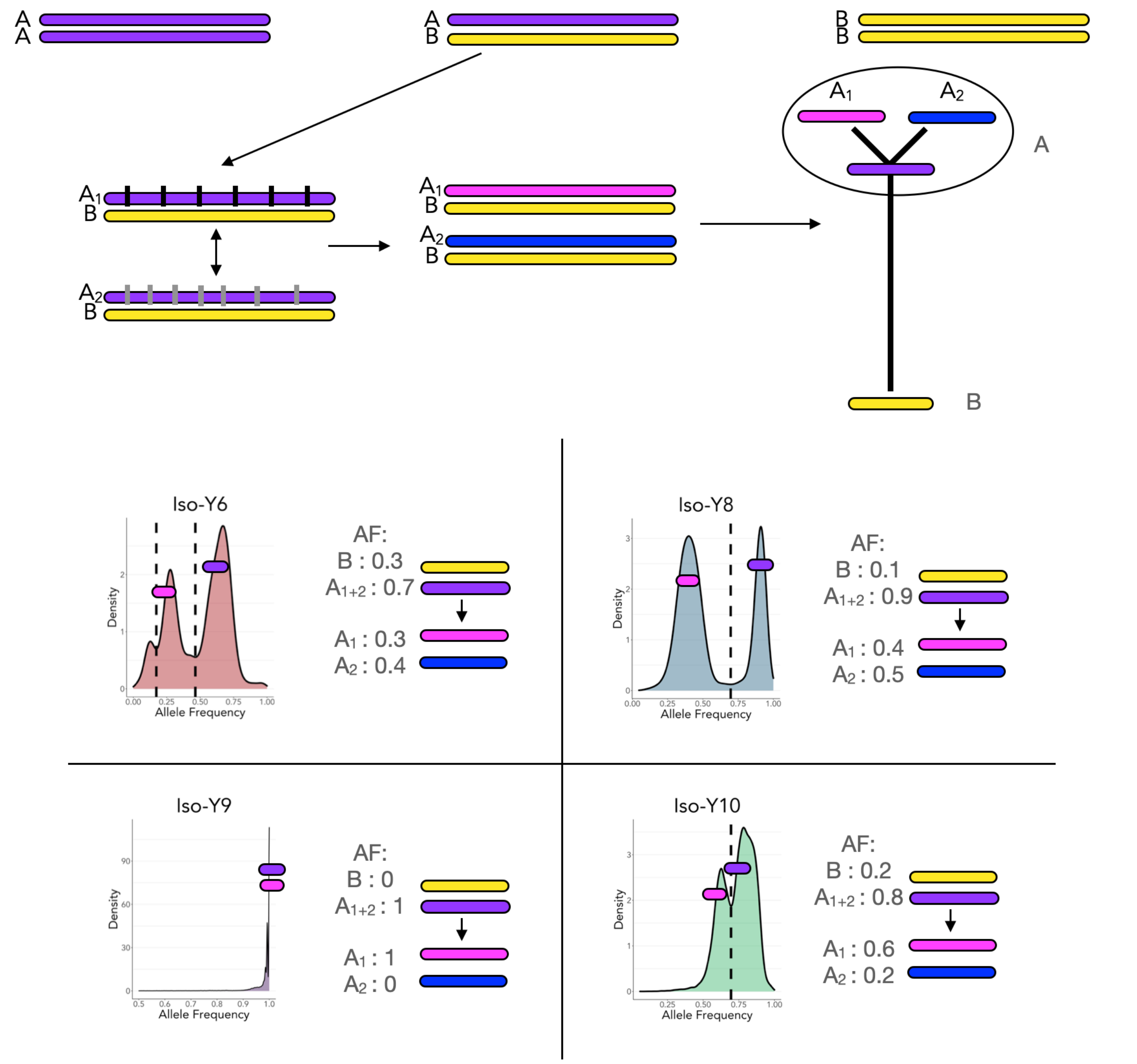


**Supplementary Figure 15.** Schematic representation of the multiple bands of allele frequencies present in the Iso-Y pool-sequencing data. AA represents homozygous, AB represents heterozygous, BB represents homozygous alternative. The ancestral A haplotype accumulates SNPs, differentiating it into two versions of the A haplotype: $A_1$ and $A_2$. The 'Y' shaped haplotype tree represents the predicted evolutionary relationships between the ancestral A, derived $A_1$ and $A_2$ haplotypes, and the B haplotype. Branch lengths represent evolutionary distance, and thus SNP count. We predict that Iso-Y9 is fixed for one of these derived A haplotypes (for illustration purposes, we have shown fixed frequencies of $A_1$). The remaining Iso-Y lines are all heterozygous with the AB genotype. There are no BB individuals. Individuals within the Iso-Y6, Iso-Y8 and Iso-Y10 pools have segregating $A_1$ and $A_2$ haplotypes, which when compared to Iso-Y9 $A_1$ show multiple bands of AFs, as shown in the calculations. Due to the nature of pool-sequencing, it is unclear what the actual genotypes are. We focus on providing an explanation of the bimodal peaks, but it is noteworthy that in Iso-Y9 there are many fixed sites, but also some "nearly fixed" sites which suggests some diversity also exists in the Iso-Y $A_1/A_1$ haplotype, which likely represents the trimodality of Iso-Y6, (i.e. further complexity in the $A_1/A_1$ haplotype that's captured in comparisons with Iso-Y6).

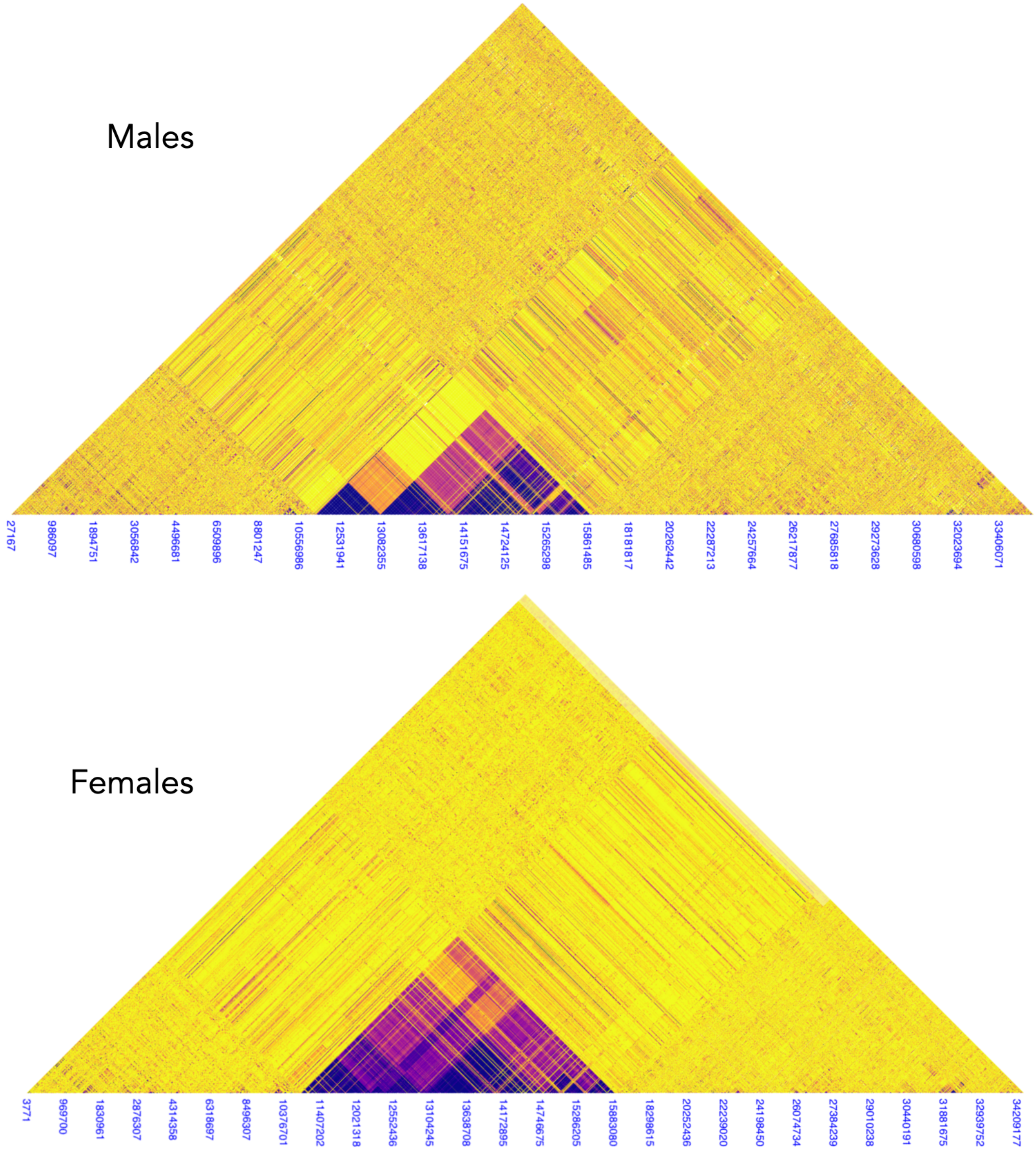


**Supplementary Figure 16.** Patterns of linkage disequilibrium (LD) in the natural data on LG1 for a) males (n=10) and b) females (n=16).

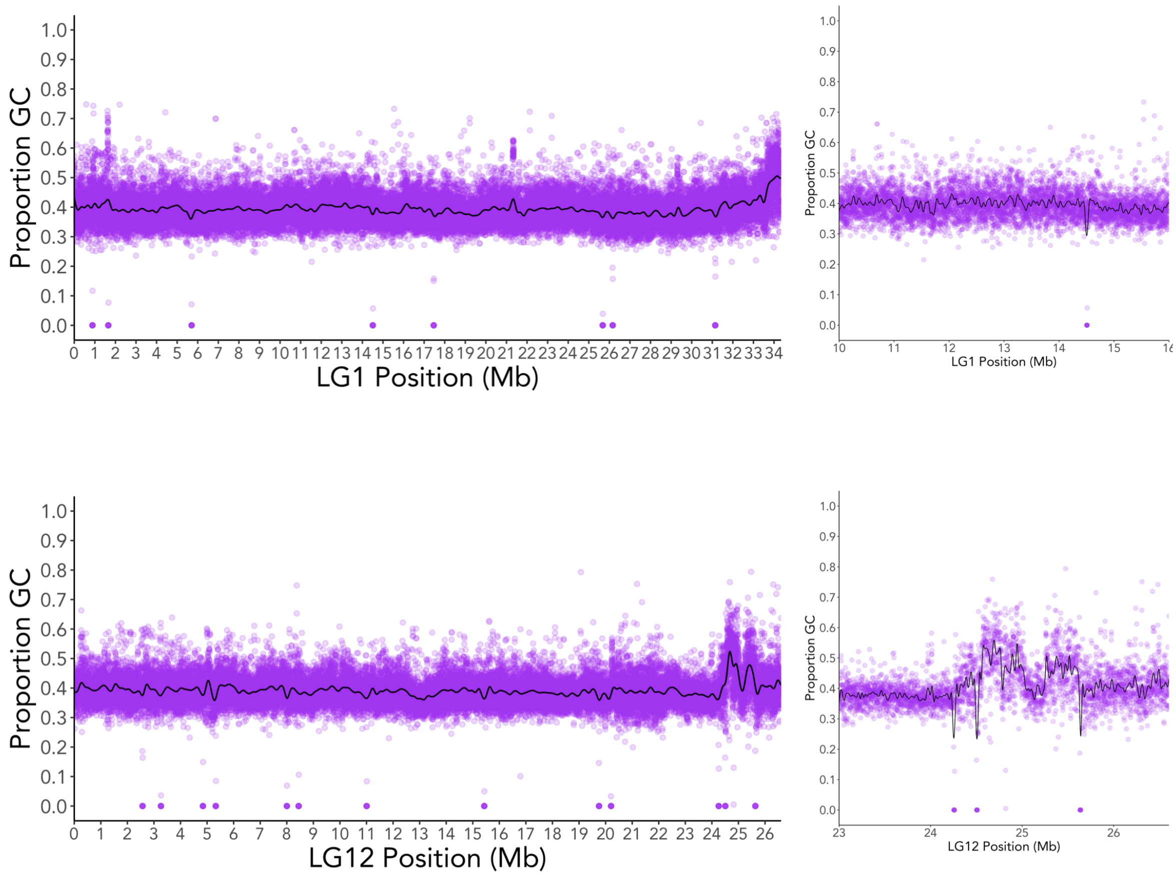


**Supplementary Figure 17.** GC content of LG1 and LG12 in 1kb windows. a) LG1 GC content; b) Zoom-in on LG1 region 2 GC content; c) LG12 GC content; d) Zoom-in on LG12 end-region GC content.

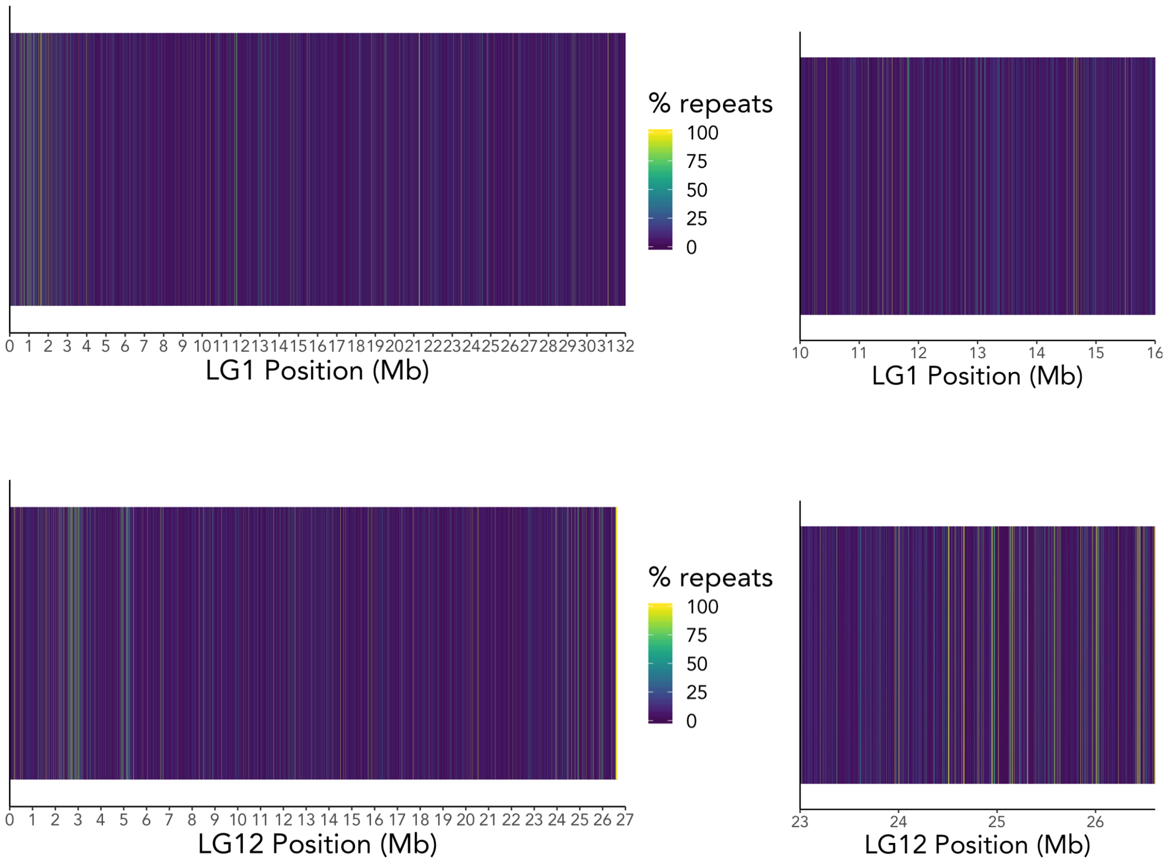


**Supplementary Figure 18.** Repeat content of LG1 and LG12 in 1kb windows. a) LG1 repeat content; b) Zoom-in on LG1 Region 2 repeat content; c) LG12 repeat content; d) Zoom-in on LG12 end-region repeat content.

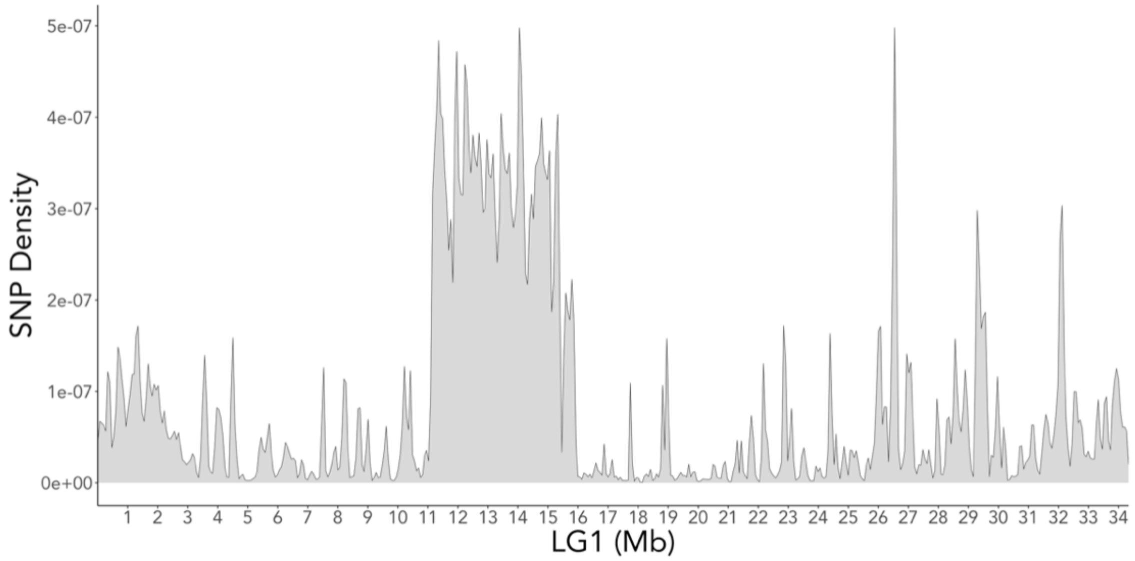


**Supplementary Figure 19.** SNP density of LG1 in the natural data (n=26).

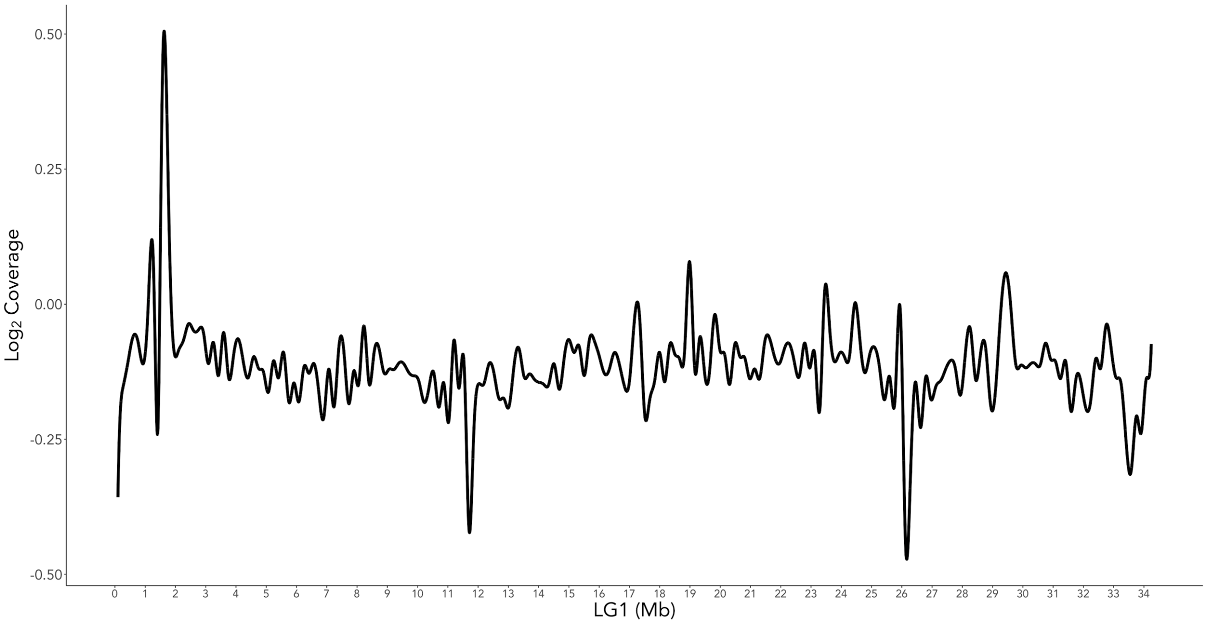


**Supplementary Figure 20.** Coverage results for the natural data.

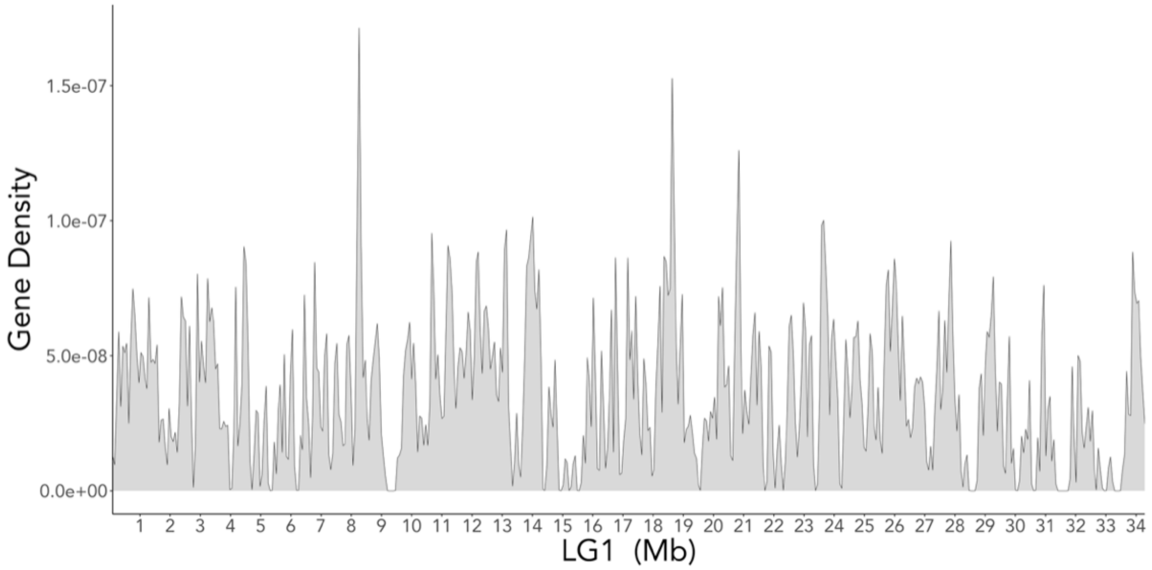


**Supplementary Figure 21.** Gene density of LG1 from the male guppy genome (ENA accession: GCA_904066995; https://github.com/bfraser-commits/guppy_genome).

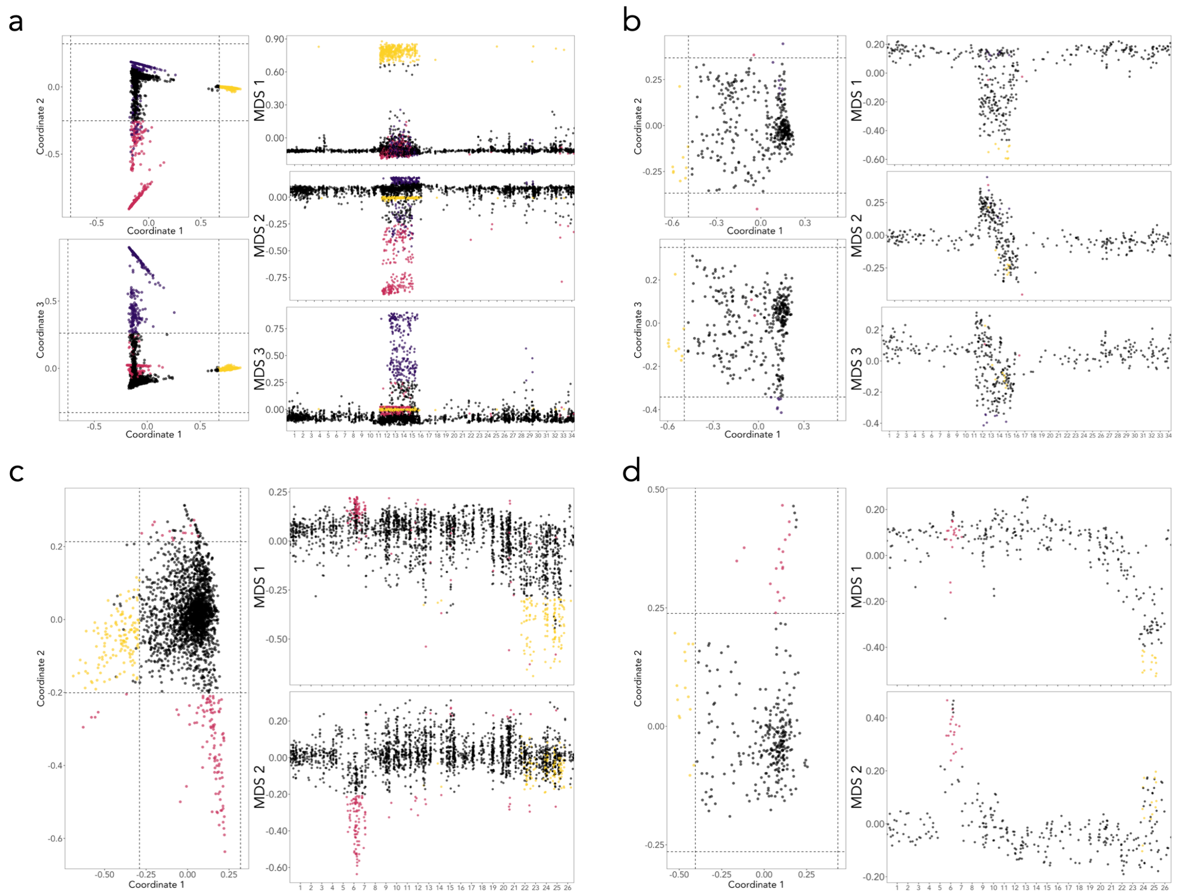

**Supplementary Figure 22.** *Lostruct* local PCA analysis for LG1 and LG12 in 10bp and 100bp windows. a) LG1 analysed in 10bp windows b) LG1 analysed in 100bp windows. LG1 plots both show the first 3 multidimensional scales (MDS: MD1: yellow; MDS2: pink; MDS3: purple). c) LG12 analysed in 10bp windows. d) LG12 analysed in 100bp windows. Although fewer windows are identified as outliers compared to 10bp windows, the same patterns of differentiation are apparent in both chromosomes.

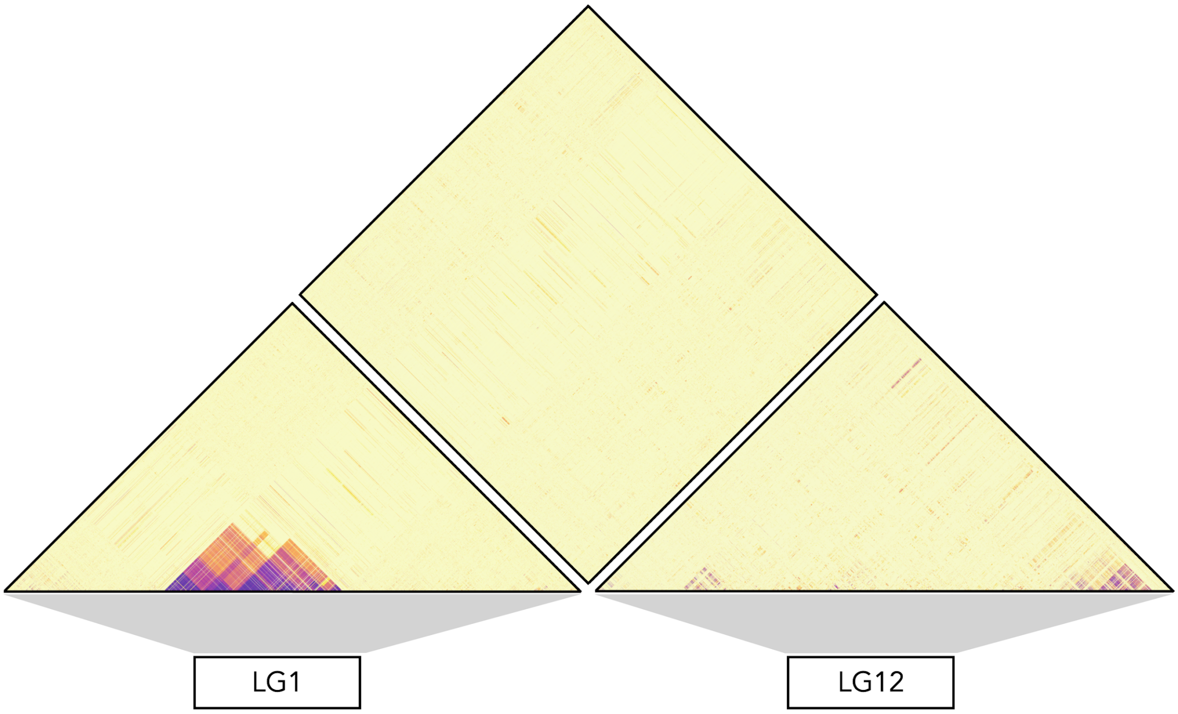


**Supplementary Figure 23.** Inter-chromosomal linkage between LG1 and LG12. Linkage information (R2) was calculated in Plink (--inter-chr), using only heterozygous sites at 5kb intervals.

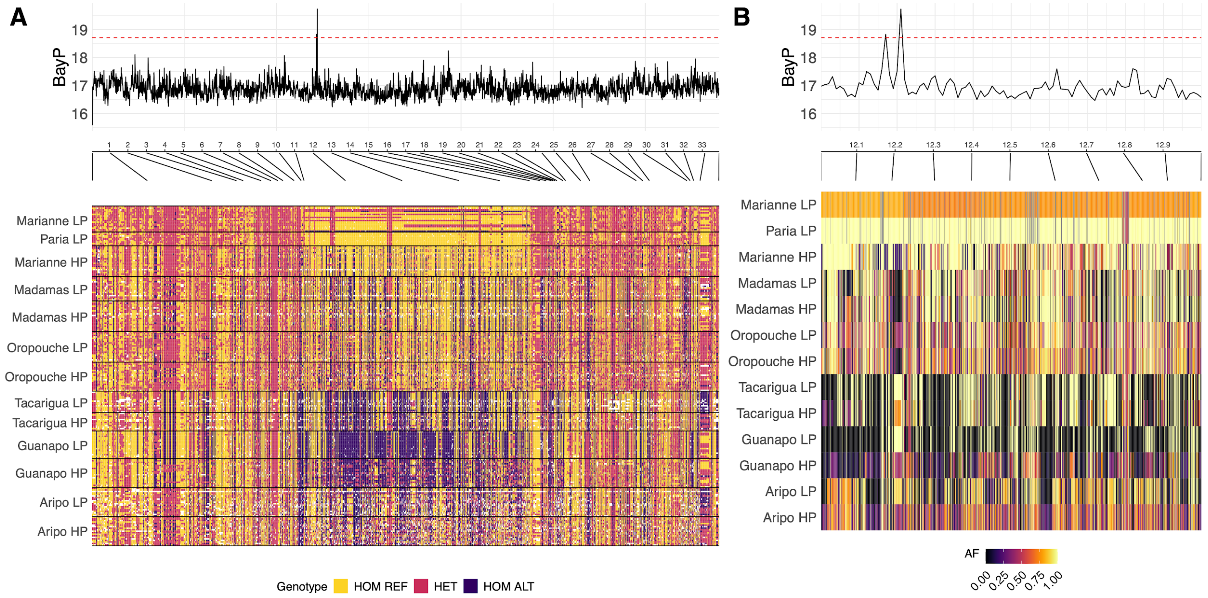


**Supplementary Figure 24.** Comparisons of LG1 among natural HP-LP populations and association with HP-LP adaptation. (a) BayPass association scores (BF) in 10kb windows along LG1, as described in Whiting et al 2020. The first two rows show genotypes of individuals from the rivers considered in this study (Marianne LP and Paria LP). Remaining populations are considered in Whiting et al 2020. Genotypes are polarised to the major allele of Paria LP. This highlights a strong peak of HP-LP association between 12 and 13 Mb that overlaps with the sex-linked region of 12.1 to 13.2 Mb described in this study. This also demonstrates that the extended haplotype structure observed across LG1 Region 2-NP is not observed across all rivers in this extended dataset. (b) shows the region between 12 Mb and 13 Mb where the signal of HP-LP association is strongest. Each row highlights the allele frequencies (AF) for each SNP within this subset of the chromosome, again polarised to the major allele in Paria LP. The strongest signal of HP-LP association here (LG1:12210000-12220000), coincides with parallel allele frequency change among all rivers, except Madamas. In this region, all LP populations (except Madamas) exhibit an increased frequency, which is sometimes fixed, relative to their corresponding HP population for the major allele that is fixed in Paria.


Whiting JR, Paris JR, van der Zee MJ, et al. (2020) Drainage-structuring of ancestral variation and a common functional pathway shape limited genomic convergence in natural high- and low-predation guppies. *bioRxiv* DOI: 10.1101/2020.10.14.339333.
