## Supplementary Text for "A large and diverse autosomal haplotype is associated with sex-linked colour polymorphism in the guppy"

**Supplementary Text 1**: π diversity across the Iso-Y lines LG1 and LG12

To explore diversity within the Iso-Y lines we calculated π across 10kb windows for our two focal chromosomes, LG1 and LG12, and performed the same multivariate approach as for Z-$F_{ST}$. Z-π-LG1 PC1 (PC1 captured 74% of the variance) represented the shared chromosomal landscape of diversity among all Iso-Y lines on LG1 and accounted for the majority of variance in diversity (Supplementary Figure 6; Supplementary Table 7). Similarly, Z-π-LG12 PC1 accounted for the majority of variance (PC1 captured 88% of the variance), and showed a significant (> critical Z-score 4.6, α=0.01) peak in diversity at 24.27 Mb (Supplementary Figure 11; SI Table 7). PC2 (PC2 captured 7% of the variance) showed a high level of residual variance in π, associated with the variance of Iso-Y9, that was not otherwise explained by the general chromosomal landscape. The diversity hotspot (peak at 24.28 Mb, > critical Z-score -1.4, α=0.01) represented by PC2 is thus unique to Iso-Y9. Both LG12 peaks: 24.27Mb (π shared by all Iso-Y lines); and 24.28Mb (π unique to Iso-Y9) overlap with previously identified male-specific contigs [1].

**Supplementary Text 2**: Examination of candidate colour and male-specific fitness genes within the identified regions of LG1

All gene annotations can be found in Supplementary Table 9. Region 1 (4-5.9Mb) contained 62 predicted genes. Of interest was *tll1*, with a role in caudal fin, dorsoventral patterning in *D. rerio* [2], *pcm1* involved in spermatogenesis [3], and protein tyrosine phosphatase non-receptor type 13 (*ptpn13*) is Y-linked in humans [4] and in some fish due to its physical linkage with *gsdf*, a gene which is highly conserved in fish sex differentiation pathways [5,6]. Region 2 (9.6-17 Mb) contained 291 predicted genes. Genes with a potential role in colour included *xpa*, involved in pigmentation and photosensitivity to UV light [7], *pcdh10a* involved in melanocyte migration, *crebbpa*, which has been identified as a candidate for plumage colouration in chickens [8], and *shoc2*, which causes pigmentation abnormalities [9], as well as five keratin genes, which have a role in pigmentation [10]. We also identified five retinal genes (*slc24a2* [11], *stra6l* [12], *pnpla6* [13], *cabp2a* [14] and *nxnl1* [15]), and three genes involved in spermatogenesis / sperm motility (*tdrd7a* [16], *nanos3* [17] and *tekt4*) [18]. Region 3 (21.9-24.9Mb) contained 94 predicted genes. This region also contained several promising candidates including two paralogs annotated as *kita*, previously identified as a key gene involved in pigment pattern formation in guppy strains [19] and zebrafish [20], *sox10a*, a sex determining region Y-box that regulates the expression of the *mitf* gene, which is the master regulator of melanophore–melanocyte differentiation in teleosts [21], and is also responsible for colouration in rock pigeons [22], *mchr1*, a melanin-concentrating hormone receptor, and a *TRYP*, located in melanocytes and involved in the production of melanin [23].

3 Sha J, Zhou Z, Li J, Yin L, Yang H, Hu G *et al.* Identification of testis development and spermatogenesis-related genes in human and mouse testes using cDNA arrays. *Mol*

*Hum Reprod* 2002; **8**: 511–517.

4 Lahn BT, Page DC. Functional coherence of the human Y chromosome. *Science* 1997.https://science.sciencemag.org/content/278/5338/675.abstract.

5 Gautier A, Le Gac F, Lareyre J-J. The gsdf gene locus harbors evolutionary conserved and clustered genes preferentially expressed in fish previtellogenic oocytes. *Gene* 2011; **472**: 7–17.

6 Liu Y, Zhang W, Du X, Zhao J, Liu X, Li X *et al.* Sexually dimorphic expression in developing and adult gonads shows an important role of gonadal soma-derived factor during sex differentiation in olive flounder (Paralichthys olivaceus). *Comp Biochem Physiol B Biochem Mol Biol* 2017; **210**: 1–8.

7 Lehmann AR, McGibbon D, Stefanini M. Xeroderma pigmentosum. *Orphanet J Rare Dis* 2011; **6**: 70.

8 Fogelholm J, Henriksen R, Höglund A, Huq N, Johnsson M, Lenz R *et al.* CREBBP and WDR 24 Identified as Candidate Genes for Quantitative Variation in Red-Brown Plumage Colouration in the Chicken. *Sci Rep* 2020; **10**: 1161.

9 Jang H, Oakley E, Forbes-Osborne M, Kesler MV, Norcross R, Morris AC *et al.* Hematopoietic and neural crest defects in zebrafish shoc2 mutants: a novel vertebrate model for Noonan-like syndrome. *Hum Mol Genet* 2019; **28**: 501–514.

10 Gu L-H, Coulombe PA. Keratin function in skin epithelia: a broadening palette with surprising shades. *Curr Opin Cell Biol* 2007; **19**: 13–23.

11 Schnetkamp PPM. The SLC24 gene family of Na+/Ca2+–K+ exchangers: From sight and smell to memory consolidation and skin pigmentation. *Mol Aspects Med* 2013; **34**: 455–464.

12 Shi Y, Obert E, Rahman B, Rohrer B, Lobo GP. The Retinol Binding Protein Receptor 2 (Rbpr2) is required for Photoreceptor Outer Segment Morphogenesis and Visual Function in Zebrafish. *Sci Rep* 2017; **7**: 16207.

13 Kmoch S, Majewski J, Ramamurthy V, Cao S, Fahiminiya S, Ren H *et al.* Mutations in PNPLA6 are linked to photoreceptor degeneration and various forms of childhood blindness. *Nat Commun* 2015; **6**: 5614.

14 Glasauer S, Neuhauss S. Expression of CaBP transcripts in retinal bipolar cells of developing and adult zebrafish. *Matters* 2016; : 1–4.

15 Cronin T, Raffelsberger W, Lee-Rivera I, Jaillard C, Niepon M-L, Kinzel B *et al.* The disruption of the rod-derived cone viability gene leads to photoreceptor dysfunction and susceptibility to oxidative stress. *Cell Death Differ* 2010; **17**: 1199–1210.

16 Tanaka T, Hosokawa M, Vagin VV, Reuter M, Hayashi E, Mochizuki AL *et al.* Tudor domain containing 7 (Tdrd7) is essential for dynamic ribonucleoprotein (RNP) remodeling of chromatoid bodies during spermatogenesis. *Proc Natl Acad Sci U S A* 2011; **108**: 10579–10584.

17 Lolicato F, Marino R, Paronetto MP, Pellegrini M, Dolci S, Geremia R *et al.* Potential role of Nanos3 in maintaining the undifferentiated spermatogonia population. *Dev Biol* 2008; **313**: 725–738.

18 Roy A, Lin Y-N, Agno JE, DeMayo FJ, Matzuk MM. Absence of tektin 4 causes

asthenozoospermia and subfertility in male mice. *FASEB J* 2007; **21**: 1013–1025.

19 Kottler VA, Fadeev A, Weigel D, Dreyer C. Pigment pattern formation in the guppy, Poecilia reticulata, involves the Kita and Csf1ra receptor tyrosine kinases. *Genetics* 2013; **194**: 631–646.

20 Parichy DM, Rawls JF, Pratt SJ, Whitfield TT, Johnson SL. Zebrafish sparse corresponds to an orthologue of c-kit and is required for the morphogenesis of a subpopulation of melanocytes, but is not essential for hematopoiesis or primordial germ cell development. *Development* 1999; **126**: 3425–3436.

21 Braasch I, Volff J-N, Schartl M. The evolution of teleost pigmentation and the fish-specific genome duplication. *J Fish Biol* 2008; **73**: 1891–1918.

22 Domyan ET, Guernsey MW, Kronenberg Z, Krishnan S, Boissy RE, Vickrey AI *et al.* Epistatic and combinatorial effects of pigmentary gene mutations in the domestic pigeon. *Curr Biol* 2014; **24**: 459–464.

23 Murisier F, Beermann F. Genetics of pigment cells: lessons from the tyrosinase gene family. *Histol Histopathol* 2006; **21**: 567–578.
